# Epidural Spinal Cord Recordings (ESRs): Sources of Artifact in Stimulation Evoked Compound Action Potentials

**DOI:** 10.1101/2024.05.17.594739

**Authors:** Ashlesha Deshmukh, Megan Settell, Kevin Cheng, Bruce Knudsen, James Trevathan, Maria LaLuzerne, Stephan Blanz, Aaron Skubal, Nishant Verma, Ben Romanauski, Meagan Brucker-Hahn, Danny Lam, Igor Lavrov, Aaron Suminski, Douglas Weber, Lee Fisher, Scott Lempka, Andrew Shoffstall, Hyunjoo Park, Erika Ross Ellison, Mingming Zhang, Kip Ludwig

## Abstract

**Introduction:** Evoked compound action potentials (ECAPs) measured using epidural spinal recordings (ESRs) during epidural spinal cord stimulation (SCS) can help elucidate fundamental mechanisms for the treatment of pain, as well as inform closed-loop control of SCS. Previous studies have used ECAPs to characterize the neural response to various neuromodulation therapies and have demonstrated that ECAPs are highly prone to multiple sources of artifact, including post-stimulus pulse capacitive artifact, electromyography (EMG) bleed-through, and motion artifact resulting from disturbance of the electrode/tissue interface during normal behavior. However, a thorough characterization has yet to be performed for how these sources of artifact may contaminate recordings within the temporal window commonly used to determine activation of A-beta fibers in a large animal model.

**Methods:** We characterized the sources of artifacts that can contaminate the recording of ECAPs in an epidural SCS swine model using the Abbott Octrode™ lead. Muscle paralytics were administered to block muscle activation preventing EMG from contaminating the recorded ECAPs. Concurrent EMG recordings of the longissimus, a long muscle of the back, were used to confirm a 2-4 millisecond (ms) latency source of EMG bleed-through that frequently contaminated the A-beta temporal window. Additionally, we obtained recordings approximately 5-10 minutes post-mortem after clear evoked A-beta and associated EMG responses ceased to characterize the representation of stimulation artifact across the array.

**Results:** Spinal ECAP recordings can be contaminated by capacitive artifact, short latency EMG from nearby long muscles of the back, and motion artifact from multiple sources. In many cases, the capacitive artifact can appear nearly identical in duration and waveshape to evoked A-beta responses. These sources of EMG can have phase shifts across the electrode array, very similar to the phase shift anticipated by propagation of an evoked A-beta fiber response across the array. This short latency EMG is often evident at currents similar to those needed to activate A-beta fibers associated with the treatment of pain. Changes in cerebrospinal fluid between the cord and dura, and motion induced during breathing created a cyclic oscillation in all evoked components of the recorded ECAP signal.

**Conclusion:** Careful controls must be implemented to accurately separate neural signal from the sources of artifact in spinal cord ECAPs. To address this, we suggest experimental procedures and associated reporting requirements necessary to disambiguate the underlying neural response from these confounds. These data are important to better understand the conceptual framework for recorded ESRs, with components such as ECAPs, EMG responses and artifacts, and have important implications for closed-loop control algorithms to account for transient motion such as postural changes and cough.

## 1. Introduction

Electrical stimulation of the spinal cord using an implanted epidural electrode array to treat chronic pain conditions was first attempted in a human subject in the 1960s (Shealy et al., 1967; Wall & Sweet, 1967). Subsequently, spinal cord stimulation (SCS) to treat multiple chronic pain syndromes has become the single largest market in the growing field of neuromodulation, with over $1 billion in sales in the U.S. annually (*Global Neuromodulation Devices Market Size Report, 2030*, n.d.). The market for SCS currently includes multiple indications, such as persistent spinal pain syndrome type 2 (PSPS Type 2), complex regional pain syndrome (CRPS), neuropathic pain, visceral abdominal pain, and intractable angina pectoris. Despite widespread use and therapeutic efficacy in many pain sufferers, many patients fail to realize significant reductions in pain and improvements in quality of life resulting from SCS (Duarte et al., 2020). This unfortunate reality exists because mechanisms of action for epidural stimulation for both the intended effect (pain reduction) and unwanted side effects (e.g., muscle contraction, paresthesia, and pain) in these indications are poorly understood.

The measurement of evoked compound action potentials (ECAPs) during stimulation-evoked epidural spinal recordings (ESRs) have been used to elucidate fundamental mechanisms of intended therapy and limiting off-target effects, as well as to provide a marker for closed-loop control of SCS (Brucker-Hahn et al., 2023; Dinsmoor et al., 2022; Lempka & Patil, 2018; Parker et al., 2012, 2013; Russo et al., 2020). Previous studies using ECAPs to understand other neuromodulation therapies have predominantly been performed in well-controlled acute settings. These studies have demonstrated that ECAP recordings are susceptible to multiple sources of artifact, including post-stimulation capacitive artifact and electromyography (EMG) bleed-through into the recordings from nearby activated muscle (Bahmer et al., 2010; Blanz et al., 2022; Miller et al., 1999; E. N. Nicolai et al., 2020; Verma, Knudsen, et al., 2023; Yoo et al., 2013). Despite these known difficulties in recording ECAPs spanning diverse stimulation targets for other therapies, a rigorous evaluation of sources of artifact in spinal cord ECAPs in a large animal model has yet to be performed.

It is well documented that the epidural surface of the spinal cord moves relative to the spinal cord during changes in posture, respiratory phase, and cough (Abejon & Feler, 2007; Friese et al., 2004; Mekhail et al., 2020; Schultz et al., 2012). Physical bodily movements can substantially influence the applied electric field at the level of the spinal cord and can cause inconsistent therapy and intermittent side effects such as motor evoked potentials, unwanted paresthesia, and pain. ECAPs have been proposed as a method to identify these changes and to provide a control signal for closed-loop SCS. However, recording electrode arrays have been shown to be very sensitive to periods of motion, which can create a large transient artifact (Giancoli, 1998; Ludwig et al., 2006, 2009; Michelson et al., 2018; Tam & Webster, 1977; Verma, Knudsen, et al., 2023; Wartzek et al., 2011). This artifact is due to the relative motion of the electrode disturbing the Helmholtz layer formed by charged molecules in tissue that can appear surprisingly similar to recorded neural signals (Bard et al., 1980; Giancoli, 1998; Merrill et al., 2005; Michelson et al., 2018; E. N. Nicolai et al., 2020; Tam & Webster, 1977; Wartzek et al., 2011). As a result, in any non-SCS studies it is common practice when conducting awake behaving electrophysiology recordings to visually identify periods of recordings contaminated by motion and remove from subsequent analysis. It is reasonable to speculate that recording ECAPs to identify movement of the electrode array during cough or postural changes happens when the electrode/electrolyte interface is also being disturbed, creating a large motion artifact which may contaminate ECAP measurements.

Prior acute neuromodulation studies used a series of controls to identify and differentiate artifacts from neural signals, but these controls have yet to be adopted into common practice for ESRs (Blanz et al., 2022; E. N. Nicolai et al., 2020; Verma, Knudsen, et al., 2023). In this study, we characterized the sources of artifacts in a swine model during epidural spinal cord stimulation using clinical SCS leads, to address the potential artifacts confounding the interpretation of ESRs. We performed a series of experimental perturbations as controls to identify the most common sources of artifact in SCS ECAP recordings including: 1) post-stimulation capacitive artifact, 2) EMG bleed-through, and 3) motion artifact induced by the cardioballistic effect associated with breathing and heartbeat. We demonstrated that recordings of high-conduction velocity A-betas are frequently contaminated by both capacitive artifact and very short latency evoked EMG, putatively from the nearby long muscle of the back (longissimus). In some cases, these sources of artifact, when viewed across the electrode array, have similar temporal latency shifts expected from evoked A-betas or A-deltas at recording sites at increasing distance from the site of stimulation. This is important because an observed temporal latency shift is often used to isolate neural components from EMG or capacitive artifact. These data are important for understanding the conceptual framework for evoked spinal cord compound action potential recordings, to understand mechanisms of action and/or optimize closed-loop systems for more consistent activation of neuronal fibers in the spinal cord.

## 2. Materials & Methods

### a. Subjects

All study procedures were approved by the University of Wisconsin—Madison Institutional Animal Care and Use Committee and were conducted under the guidelines of the American Association for Laboratory Animal Science in accordance with the National Institutes of Health Guidelines for Animal Research (Guide for the Care and Use of Laboratory Animals). The swine model was selected as the size of the swine spinal cord, including the organization of the roots/rootlets as they enter the cord at a given spinal level, have been shown to be highly similar to the human (Busscher et al., 2010; Cuellar et al., 2017; Schomberg et al., 2017; Sheng et al., 2010). Subjects included seven healthy domestic (Yorkshire/Landrace crossbreed) swine. One subject from the cohort was excluded from data analysis due to complications during the surgical procedure precluding a complete set of accurate spinal cord recordings (analyzed cohort 4F/2M; mean ± SD = 33.3 ± 3.8 kg). All subjects were housed individually (21◦C and 45% humidity) with *ad libitum* access to water and were fed twice daily. Each subject was given an intramuscular, injectable induction anesthesia cocktail of telazol (6 mg/kg) and xylazine (2 mg/kg). Temperature, capnography, pulse rate and EKG were recorded via AD Instruments PowerLab 8/35 and visualized in real time via LabChart (ADInstruments, Sydney, Australia). An intravenous catheter was placed in a peripheral ear vein for drug and fluid administration. Subjects were endotracheally intubated and maintained with a mechanical ventilator using 1.5–3% isoflurane. All vital signs including temperature, heart rate, CO_2_, and respiration were continuously collected and recorded every 10 minutes and used to monitor depth of anesthesia.

### b. Surgical & Imaging Methods

In a ventral recumbent position, a dorsal midline incision was made to allow for a laminectomy, from the L1 to L3 spinal levels. The bone and connective tissue, with dura intact, were removed from between spinal processes to create an insertion window, such that there was consistently a window over the stimulating contacts. A window was then created at each spinal level to visualize the implantation of the two 8-contact, platinum iridium, Octrode™ leads (Abbott Neuromodulation, Plano, TX). Utilizing fluoroscopy and partial laminectomies to visualize and optimize the location of electrodes (T10-T15), the two Octrode™ leads (8 contacts each) were placed sequentially. One electrode array (contacts 1-8) was placed caudally, and one array (contacts 9-16) was placed cranially over the midline, and in such a way as to prevent any overlap of the contacts (Figure 1A and B). The two most caudal electrode contacts (7 and 8) were used as stimulation electrodes with the other 14 electrodes used for ESRs. Each contact was 1.4 mm in diameter, 3 mm in length, with a 7 mm center-to-center distance separation between adjacent contacts (Figure 1B) Leads were advanced until all 16 contacts were at the approximate midline, on the dura, and spanned the T10 to L1 region of the spine. Fluoroscopy was used to confirm location of all electrode contacts. Across all subjects, stimulating contacts (contacts 7 & 8) spanned the T14/T15 level (n=2), the T15/L1 level (n=3), and the L2/L3 level (n=2) (Figure 1C).

**Figure 1.**
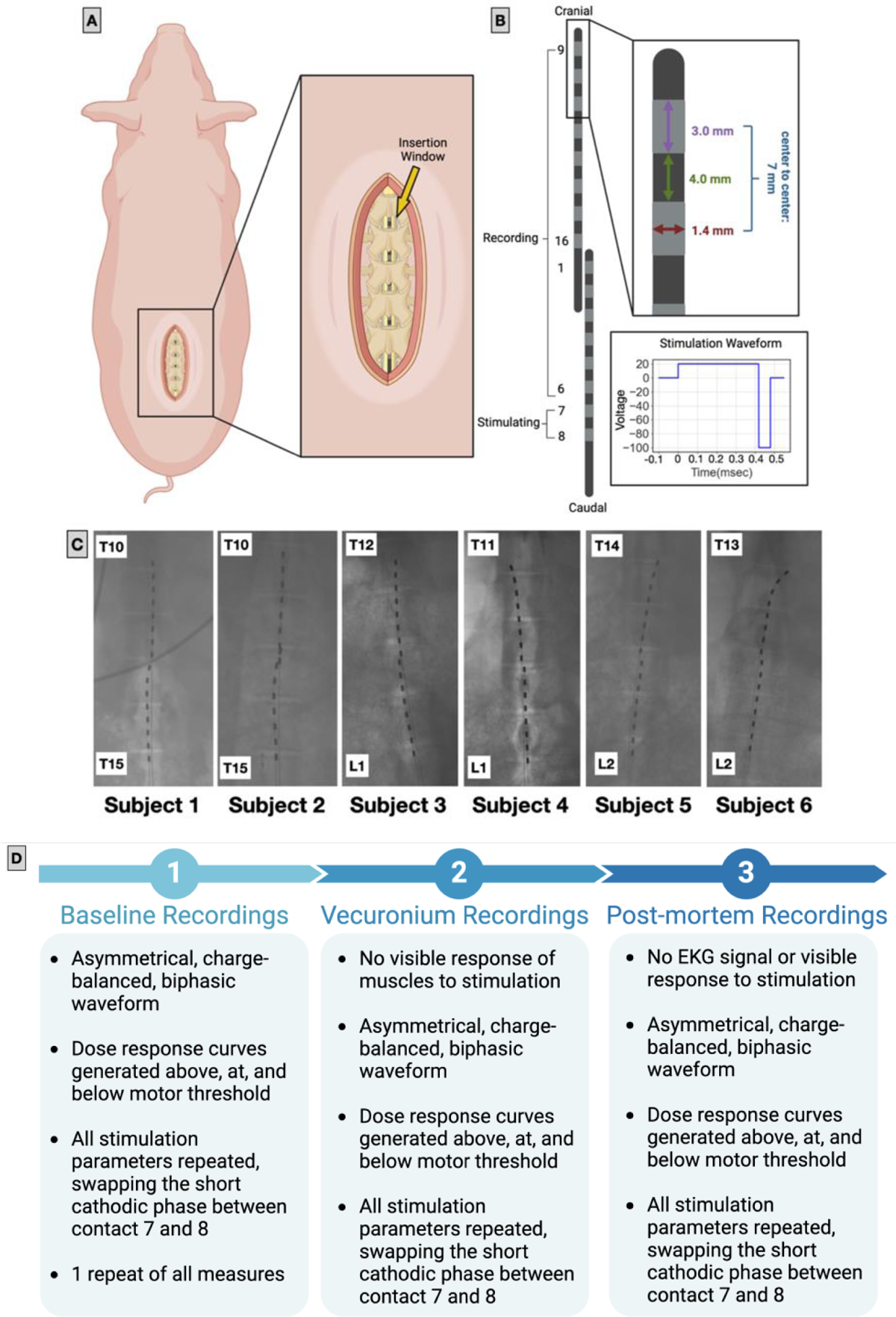
Experiment Overview: (A) Schematic of swine spinal cord cut down location, and representative positioning of the epidural electrode arrays, with surgical insertion windows (inset). (B) Schematic of the Abbott Octrode™, with stimulating contacts (7 & 8) located at the most caudal end of the caudal lead, and the remaining 14 as recording contacts. The diameter of each electrode contract was 1.4 mm, and length 3 mm; spacing between contacts was 4 mm and 7 mm center to center. The asymmetric biphasic stimulation waveform used to determine effects of activation on the spinal cord (C) Fluoroscopy images of the placement of electrodes in all subjects with annotated spinal levels (n=6). (D) Experimental Timeline: Baseline recordings were obtained following the determination of visual motor thresholds (amplitude), following baseline recordings Vecuronium (skeletal muscle paralytic) was administered and baseline stimulating parameters repeated. Following euthanasia, post-mortem recordings were collected with all baseline stimulation parameters.

Variations between stimulating location (Figure 1C) were due to landmark error, as location was based on the presence of the most caudal rib, which was difficult to visualize on CT in some cases. Additionally, in domestic swine the number of thoracic vertebrae is not fixed, and can vary between 15 and 17 thoracic segments (Zhang et al., 2015). All final placements were confirmed during post-mortem microdissection with L1 being designated as the segment with the most caudal rib. Once electrodes were positioned, the incision was covered with sterile saline soaked gauze to prevent drying of the surgical pocket throughout the experiment.

To monitor EMG activity from surrounding back muscles, innervated by the L1/L2 region, bipolar stainless steel fine wires (200 mm long, 0.002 inch diameter) with enamel insulation, and exposed tips (2 mm), were placed in the muscle using a needle (Rhythmlink, Columbia, SC). Additional in-house made EMG electrodes were used for comparison to the prefabricated electrodes (data not shown). These electrodes were produced using a 7-strand, insulated, stainless steel wire, with insulation removed from the tips (A-M Systems, Sequim, WA). EMG electrodes were initially placed (subjects 1 and 2) in the lower limb muscles (semitendinosus, medial gastrocnemius, tibialis anterior). However, following two consecutive experiments with little to no EMG response to stimulation in these instrumented muscle groups, microdissection revealed one of the primary muscles innervated at the T15/L1 level to be the longissimus (Supplemental Figure 1). The remaining five subjects (subjects 3-7) underwent instrumentation of the longissimus. All EMG recordings were collected using a differential recording configuration with a 25 kHz sampling rate.

### c. Stimulation & Recording

The Tucker-Davis Technologies (TDT) RZ2 electrophysiology system (Alachua, FL) was used to control stimulation and to record both EMG and ECAP signals. The stimulating leads were connected to the TDT system with a multi-lead trial cable adapter, to a Threshold NeuroDiagnostics Medusa cable (Threshold NeuroDiagnositcs, New York, US). The spinal cord ECAP signals were sampled at ∼25 kHz. A local reference was placed in adipose tissue to prevent the differential subtraction of ECAP components which can occur when differential recordings are measured using a separate electrode contact as a reference. This occurs because both the reference and recording electrode will measure the evoked action potential as it propagates down the array. Depending on the relationship of the reference electrode to the stimulating sites, this will impact changes in latencies recorded at different sites along epidural lead (Verma, Romanauski, et al., 2023). As the reference was placed near the cord in adipose tissue, it should still function to mitigate common source of artifact (e.g., EMG bleed-through, stimulation artifact) without also differentially subtracting intended ECAP components.

At the beginning of all experiments the visual motor threshold was determined by administering low amplitude stimulation, then increasing stimulation amplitude until there were visible A-beta signal recordings, and visible constriction/activation of innervated muscles (longissimus). Once threshold was determined, a protocol of stimulation amplitudes below, at, and above threshold were selected and randomized throughout the remainder of the experiment to avoid any cumulative effects of low to high stimulation. Dose-response curves were generated using multiple stimulating and recording configurations in each animal to assess the impact of each configuration on the stimulation artifact. Across all subjects, stimulation consisted of a charge-balanced waveform at 42 Hz and varying amplitudes. 42 Hz was selected as it is not a factor of 60 Hz noise, such as 20 or 30 Hz, and therefore 60 Hz noise can be attenuated by averaging over multiple ECAP presentations. For all ECAPs, asymmetric biphasic pulses with a long 400 µs phase followed by a short 80 µs phase were used to minimize deviation from waveforms, commonly used for SCS, while simultaneously minimizing contamination of ECAPs with capacitive artifact.

Based on chronaxie curves, established for perception threshold associated with SCS in humans, a first phase, that is 1/5th the amplitude and 5 times longer than the second phase, should not activate until the amplitude of the first phase reaches ∼2.5x the activation threshold associated with the second high-amplitude phase of the waveform **(Ranck, 1975; Yearwood et al., 2010)**. Additionally, anodic stimulation is expected to have a **∼3x larger threshold for activation of the same tissue** (Ranck, 1975). This optimization of the waveform attempts to minimize the capacitive stim artifact and also cause nerve activation to occur near the end of the first phase or during the second phase of the asymmetric waveform. This maximizes separation of ECAP signals from the stimulation artifact for the best differentiation of neural signals. Selecting language based on the phase during which neural activation was expected to occur, henceforth: 1) Stimulation ‘amplitude’ refers to the amplitude of the short-duration, high-amplitude, second phase of the biphasic waveform, and 2) **‘Anode’/’Cathode’, refer to the electrode contact, and the associated direction of current flow applied during the short-duration high-amplitude phase of the asymmetric waveform.**

Following the baseline stimulation paradigm, vecuronium was administered to paralyze the muscles, eliminating any potential EMG signal contaminating the recorded ECAP of the spinal cord. Following experimental procedures, the isoflurane inhalant was increased to 5% and animals were euthanized with a super saturated KCl solution delivered intravenously at 1 mL per kg. Post-mortem recordings were verified by loss of heart rate signal, loss of evoked EMG responses, and loss of ECAP responses at distal recording sites minimally contaminated by capacitive stimulation artifact. Post-mortem recordings were taken to identify biological signal and sources of noise, stimulation artifact, capacitive decay of the stimulation artifact, and instrumentation-related sources of noise.

### d. Data Analysis

The pyeCAP package (https://github.com/ludwig-lab/pyeCAP, version 0.0.1) was employed in Python 3.7 for offline analysis of electrophysiology data. The adopted filtering strategy aimed at applying the minimum necessary filtering to mitigate the influence of prevalent noise sources without introducing notches at 60, 120, and 180 Hz to reduce powerline (60 Hz) noise. Subsequently, a median high pass filter, spanning 201 samples (8.2 ms) and a Gaussian low pass filter with a cutoff frequency of 5.2 kHz were applied. The median high-pass filter, serving as a non-linear filter, removes signal drift over time durations significantly longer than the signals of interest, avoiding any filter ringing caused by the high-frequency and high-amplitude components of the stimulation artifact (Jarske & Vainio, 1993). Importantly, the averaging of repeated ECAPs effectively reduces noise across the frequency spectrum without inducing ringing. In this context, white noise is diminished by a factor of √*N*, where *N* represents the number of stimulation pulses, totaling 240, resulting in an over 15-fold reduction in white noise amplitude from the averaging of repeated signals alone. Given this averaging, the influence of high-frequency noise on the signals of interest was minimal, allowing for a higher low-pass filter cutoff frequency of 5.2 kHz. This cutoff is substantially above the frequency content of our signals of interest, and the Gaussian filter’s slow roll-off is crucial to minimize any ringing effects from the high frequency and amplitude components of the stimulation artifact (Widmann & Schröger, 2012). Consequently, the Gaussian low-pass filter’s impact on the final eCAP response was negligible and could be entirely omitted with minimal effect on the analysis. Our preprocessing strategy, focused on preserving the integrity of the recorded eCAP signal, addresses the potential for filter ringing around stimulation pulses to produce artifacts with temporal characteristics similar to neural signals—a critical consideration often overlooked in filtering strategies aimed solely at noise reduction (Widmann & Schröger, 2012). ECAP component amplitude was quantified as the area under the curve (AUC) using the root mean squared method (RMS) calculated over the time period from 1-2 ms following stimulation onset, effectively encompassing the time window during which the A-beta activity is expected to occur (Blanz et al., 2023; Haughton et al., 1994; Kent-Braun, 1999; Zory et al., 2005). Similarly, the EMG component amplitude was also calculated as the RMS amplitude, but from 4-10 ms after the stimulus pulse.

An additional analysis of how physiological processes such as breathing, and heart rate could modulate ECAP signals was performed by first extracting the RMS amplitude of A-beta and EMG signals following each stimulation pulse. Then, to analyze the influence of low-frequency physiological phenomena, we utilized a zoom-fast Fourier transform (zoom-FFT) technique (Hoyer & Stork, 1977). The zoom-FFT method is advantageous for its ability to focus on a narrow frequency range—in this case, 0-2 Hz—thereby providing enhanced resolution of spectral components within this specific bandwidth. This is achieved by digitally filtering the signal around the frequencies of interest, in this case with a Butterworth filter, selected for its flat frequency response in the passband to minimize distortion of the low-frequency signals. Subsequently the data was padded to increase the frequency resolution of the FFT, following the application of which only frequencies below the filter cutoff were retained. This effectively magnifies the spectral detail of the bandwidth of interest without the loss of resolution associated with a broad-spectrum FFT analysis. For the statistical validation of our findings within this narrow frequency band, we implemented a bootstrap approach to generate a 95 percent confidence interval for the FFT spectrum. This was performed by randomly reshuffling the signal data 1,000 times and recalculating the FFT for each shuffle. This method allows for the estimation of the variability and significance of peaks within the FFT, distinguishing them from the baseline noise level with statistical confidence. By comparing the original FFT spectrum to the distribution obtained from the bootstrap samples, we were able to identify frequencies that consistently stood out from random variations, suggesting a possible physiological origin. All analysis performed on this data was conducted *post-hoc* and should be considered exploratory.

## 3. Results

Current closed-loop paradigms for SCS are often predicated on the concept of ‘gate theory’ (Melzack & Wall, 1965). This theory suggests large diameter (low activation threshold) sensory A-betas fibers within the dorsal column and/or in the root/rootlets are activated to inhibit the input signals from (high activation threshold) C-fibers associated with chronic pain via multiple hypothesized pathways. In this paradigm, the goal is to generate consistent activation of the A-beta fibers to help desensitize the central nervous system to all inputs from the same dermatome. Activation of A-delta fibers associated with acute pain, and the direct/indirect^1^ activation of A-alpha motor fibers are considered off-target effects that ultimately limit the ability to maximally activate intended A-betas. Consequently, SCS ECAPs are commonly used to identify the activation of A-beta fibers that transmit action potentials at relatively high conduction velocities (30-70 m/s) (Erlanger & Gasser, 1930; Gasser, 1941, 1960; Parker et al., 2012), while avoiding stimulation current amplitudes that also evoke slower ECAP components. These slower ECAP components suggest activation of unwanted/off-target nociceptive A-delta fibers or muscle activation.

We evaluated each component of the SCS ECAP with a particular focus on the epoch in which recorded signals from A-beta fibers are anticipated to appear. Initial visual inspection suggested that the A-beta window may be contaminated by a very short-latency EMG signal and substantial post-stimulation capacitive artifact. To positively confirm the sources of these artifacts, we first administered a muscle paralytic to positively confirm the source of bleed-through was indeed EMG, and that the paralytic was of sufficient dose to dramatically reduce or eliminate observed direct EMG recordings. We then compared the relative timing of EMG from the long muscles of the back, the most obvious short pathway for an SCS evoked muscle response, to putative EMG bleed-through in the SCS ECAP. Third, recordings were made approximately 5-10 minutes after euthanasia to assess the extent of stimulation artifact contamination into the A-beta window. The 5-to-10-minute waiting period was selected as a trade-off between the minimum time needed to eliminate neural components of the SCS ECAP or EMG response, and deterioration of the prep post-mortem, no longer representing the *in-vivo* condition. Finally, evoked SCS ECAPs were analyzed post-hoc, and compared to respiration/cardiac cycles, periods of tremor, and large stimulation evoked muscle responses. This analysis was performed with the goal of directly comparing periods of ECAP recordings with no movement to periods of ECAP recordings with visually identified periods of movement that may induce relative motion of the electrode arrays.

### a. ECAP Recordings Contain Multiple Components

Initial visual inspection of recorded ECAPs across the array revealed multiple components, including the stimulation artifact, A-beta fiber signals, and EMG contamination from surrounding muscles. Figure 2 depicts a representative pulse-by-pulse and averaged stimulation evoked ECAP illustrating the multi-component signal (stimulation: 480 µs pulse width, 240 pulses, 42 Hz, 7 mA). Even short-latency ECAP signals, such as those generated by the activation of high conduction velocity A-beta fibers, can be contaminated by both capacitive artifact and very short-latency evoked EMG, putatively from the nearby long muscles of the back (Figure 2). It was also noted that the non-averaged pulse-by-pulse ECAP traces frequently had outlier traces or even a bimodal distribution of traces (Supplemental Figure 2). This grouping of outliers (indicated by the blue arrowheads) may be suggestive of neural ECAPs contaminated by motion artifact and is analyzed in a subsequent section (Results: Section G).

**Figure 2.**
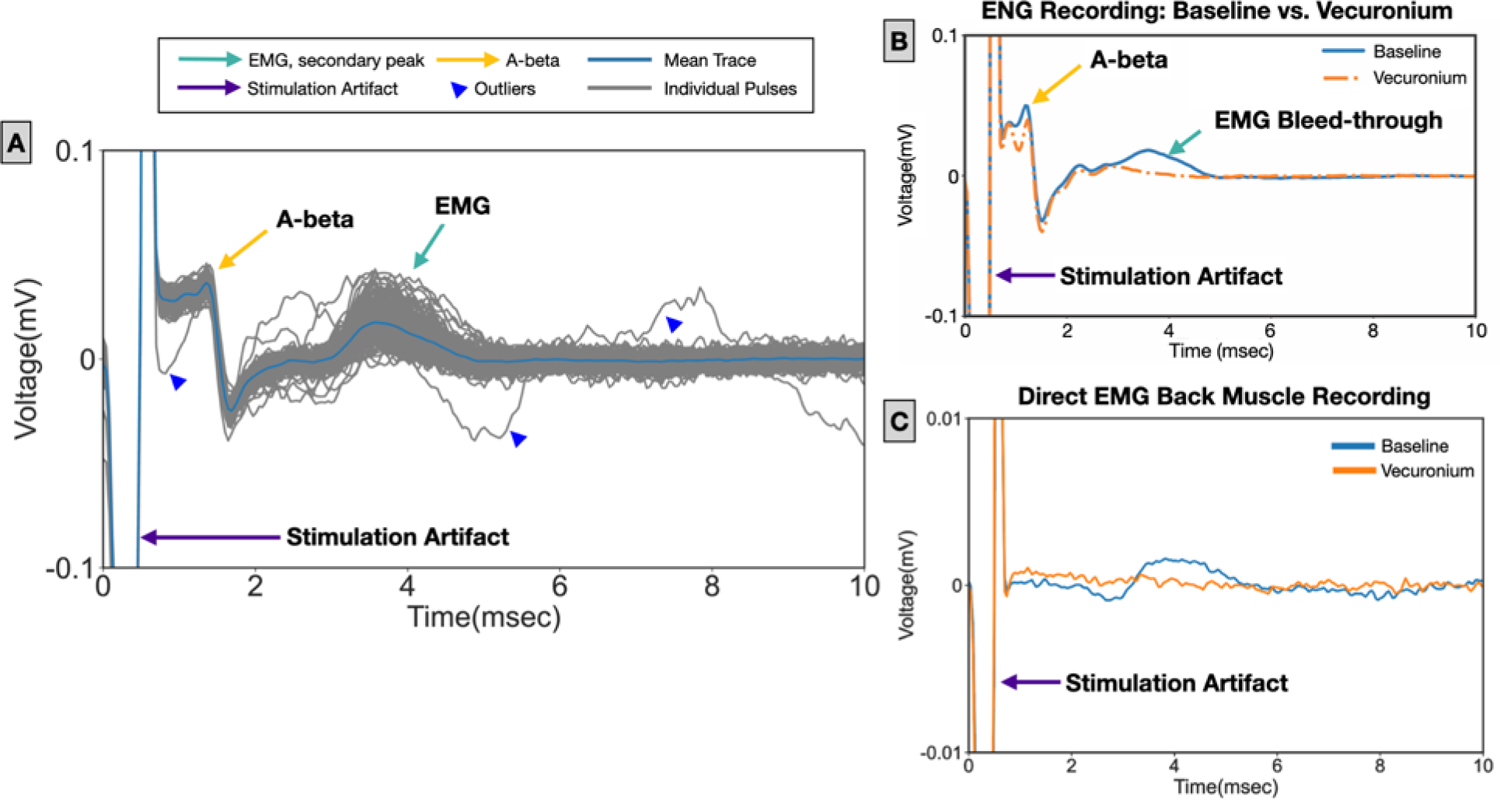
**(A)** Representative (Subject 6) pulse-by-pulse (gray lines) and mean (blue line) ECAP recordings following stimulation (7 mA, short cathodic pulse at contact 7). ECAP traces contain the stimulation artifact (purple arrows), A-beta fiber response (yellow arrows), and EMG signal (green arrows). Blue arrowheads indicate the grouping of non-gaussian outliers, putatively caused by motion artifact. **(B)** Mean ECAP recordings during baseline (blue line) and under vecuronium (orange). **(C)** Direct EMG recording of the back muscle (mean) during baseline stimulation (blue) and under vecuronium (orange).

### b. Hypothesized EMG Components Eliminated After Administration of Vecuronium

Vecuronium, a skeletal muscle paralytic, was administered following baseline recordings. Paralytic and baseline recordings were compared at the same stimulation doses to separate EMG bleed-through from neural signal and stimulation-evoked capacitive artifact (Figure 2B and C). As hypothesized, the putative short latency muscle responses were eliminated from the ECAPs after the administration of vecuronium. The disappearance of the EMG signal confirmed that these components of the response were due to the electrical signal generated by the activation of muscles close enough to the spinal cord to bleed into the ECAP recordings (Figure 2B). After verifying the short latency response was indeed EMG bleed-through in early animals, post-mortem anatomical microdissection was used to identify the longissimus muscle as having one of the shortest lengths of nerve before termination emerging from the T15/L1 levels. Direct EMG recordings from the longissimus muscles taken in later animals confirmed a consistent evoked muscle response that began between 2-4 ms after each stimulation artifact (Figure 2C, and Supplemental Figure 3 for cohort analysis).

### c. ECAP Components Across the Array

In Figure 3 the averaged ECAP recordings across the array from contacts 1-6, and 9-16, are shown at an amplitude where each component had the least overlap for one animal. The short activating cathodic pulse (cathode), which was expected to have the lowest threshold for fiber activation compared to the short anodic pulse (anode), was initially placed at the more cranial contact of the bipolar pair (contact 7) to avoid the electric field created by the anode (contact 8) from interfering with the propagation of the evoked action potentials generated by the cathode. On all contacts, at time t = 0 ms, there is a stimulation artifact that has limited pulse-by-pulse variability. As expected, the representation of the stimulus artifact changes dramatically based on the distance of the recording site from the stimulating bipolar contacts. The magnitude of the stimulation artifact during the period of stimulation changed based on distance and, as the initial visual inspection suggested, there was a presumed capacitive decay component lasting well after the stimulation period that became both larger in amplitude and longer in duration at the recording sites closest to the bipolar pair (Figure 3, blue opaque region). The minimum distance required to separate the A-beta component from the stimulation artifact was contact 1, or the last recording contact on the caudal array. This would indicate that for a reliable A-beta signal, a separation distance of at least 35 mm or more is required from the stimulation contacts to the recording contacts for stimulation amplitudes required to generate ECAPs.

**Figure 3.**
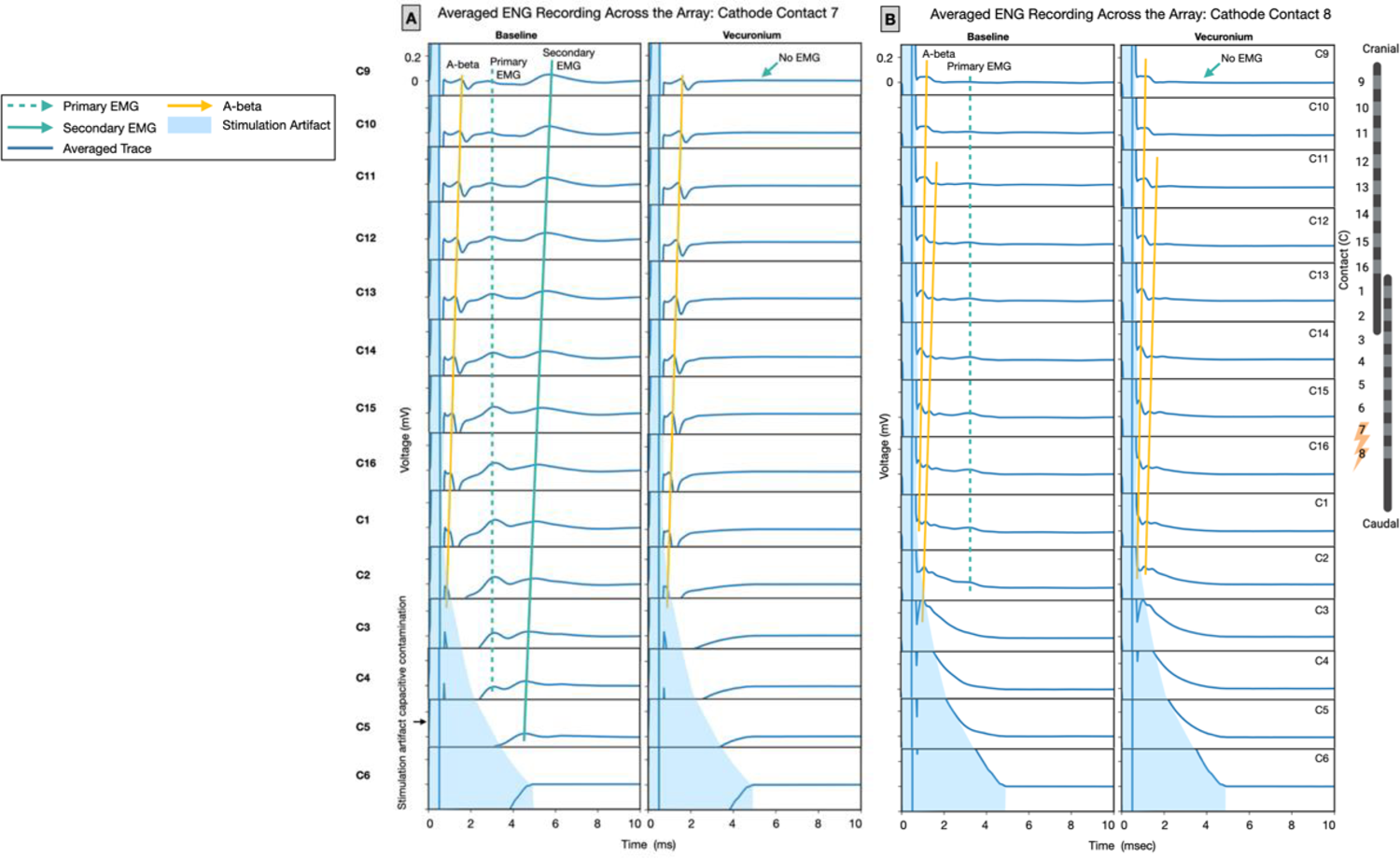
Representative propagation of A-beta and ECAP components across the array in one subject (Subject 4). **(A)** Stimulation with cathode at contact 7, the mean ECAP recordings are shown at baseline (panel 1) and after administration of vecuronium (panel 2). **(B)** Stimulation with cathode at contact 8, ECAP recordings are shown at baseline (panel 1) and after vecuronium (panel 2). A-beta signals are indicated with yellow, solid lines. The primary EMG signal is indicated with the dashed green line, and secondary EMG with the solid green line (Gmel et al., 2023). The region of stimulation artifact is noted by the opaque blue region. Each row represents one of the fourteen recording contacts (C1-14), with the top rows being furthest from, and bottom the closest to, the stimulating contacts (Supplemental Figure 4 for all subjects). Note the secondary EMG component denoted by the solid green line is eliminated by vecuronium, but still appears to have a ‘phase propagation’ across the recording array that would be consistent with neural evoked activity. Consequently, depending solely on the appearance of phase propagation across the array would be insufficient to identify whether recording components are neural or muscular.

At recording sites furthest from the stimulation electrodes, there was a noticeable evoked signal approximately 1-2 ms after the stimulation pulse (Figure 3, yellow line). This feature is most consistent with what prior studies have identified as the neural signal generated by activation of A-beta fibers. Based on the latency of this response, estimation of the conduction velocity was in the 30-70 m/s range, also consistent with the expected conduction velocity of A-beta fibers (Erlanger & Gasser, 1930; Gasser, 1941, 1960; Manzano et al., 2008; Parker et al., 2012). As the recording sites were sampled across the array, closer to the stimulating pair, the latency of this feature decreased, as expected, given the evoked action potential should arrive at closer recording sites more quickly. The amplitude of this recorded evoked response tended to decrease slightly and becomes more diffuse as distance increased from the stimulating contacts, consistent with previous observations of the behavior of ECAPs based on recording distance (Andreis et al., 2021; Verma, Knudsen, et al., 2023). At recording sites progressively closer to the stimulating pair (contacts 3-6) this evoked feature moved visibly closer to the stimulation artifact until partially overlapped, and for the closest recording sites, was no longer visible.

At higher stimulation amplitudes two additional components were often visible in the evoked compound action potential recording, one typically spanning 2-4 ms post-stimulation and another spanning 6-10 ms post-stimulation. Notably, the shorter latency feature could overlap with the putative A-beta response, depending on the distance between stimulation and recording electrodes. These features were qualitatively consistent with our prior experience in evaluating EMG bleed-through into ECAP recordings evaluated by our lab when stimulating and recording from the vagus nerve in large animal models (Blanz et al., 2022; E. N. Nicolai et al., 2020). These data, along with the short latency of the response suggested a localized muscle response close to the location of the array on the spinal cord.

### d. EMG Bleed-through was Highly Dependent on Electrode Recording Location

EMG bleed-through was highly dependent on electrode recording location (Figure 3, solid and dashed green lines). Importantly, in some cases, the EMG component shifted phase across the recording contacts (Figure 3A Baseline, Secondary EMG). At times this could give the false appearance of the change in latency expected from an evoked action potential progressively propagated to more distal recording sites, especially if only tracked across a limited subset of the electrodes (Figure 3). The latency shift of the EMG across recording sites could frequently appear very similar to slower ECAP components like A-deltas or a synaptic delay. However, this signal is eliminated by the administration of a muscle paralytic (vecuronium), confirming it as a muscle response (Figure 3A and B, Vecuronium).

### e. Post-mortem Stimulation Confirmed Neural Signal in ESRs

In our previous work we have confirmed neural recordings by transecting the peripheral nerve of interest (Blanz et al., 2022; E. N. Nicolai et al., 2020). In this study, recordings were taken after the animal was sacrificed (t=5 to 10 minutes post-mortem) to eliminate the stimulation evoked neural component, which allows for identification of stimulation evoked capacitive component of the stimulation artifact which may be conflated with neural response. Prior studies have demonstrated that CNS neurons cease their activity between 5 and 15 minutes post euthanasia if not excised and transferred to media containing oxygen/nutrients (Heymans, 1950). Therefore, stimulation dose-response curves were repeated, and recordings were taken 5 to 10 minutes after euthanizing to eliminate recorded biological responses and identify and characterize stimulation artifact capacitive components that contaminate the recording window beyond the period of stimulation. As it is possible to have some stimulation evoked activity for a short period after termination, loss of clear A-beta SCS ECAPs components and EMG of the back muscle were used to confirm the full loss of stimulation responsiveness. To positively confirm full loss of neural activity, both putative A-beta and back muscle EMG responses observed in the live SCS ECAP recordings at the highest dose administered needed to be eliminated.

Figure 4 shows a representative direct comparison of pre-mortem and post-mortem ECAP recordings at the subject-specific maximum amplitude, at a recording location suspected to have substantial capacitive artifact contaminating the A-beta region of recordings. In the pre-mortem baseline condition several neural components can be seen, which at the most distant recording sites are separate and distinct from the capacitive portion of the stimulation artifact. At closer sites, these components are partially superimposed on the capacitive artifact, and are eliminated in the post-mortem recordings leaving only the capacitive artifact. While the size of this post-stimulus capacitive artifact could change between pre- and post-mortem conditions, due to changes in tissue properties, this data clearly shows the substantial capacitive artifact that overlaps and contaminates the ECAP signal (Figure 4). As the real A-beta is riding on top of the capacitive artifact, the summation of the waveforms can appear very different than the purely capacitive artifact post-mortem after the ‘neural’ component has been eliminated.

**Figure 4.**
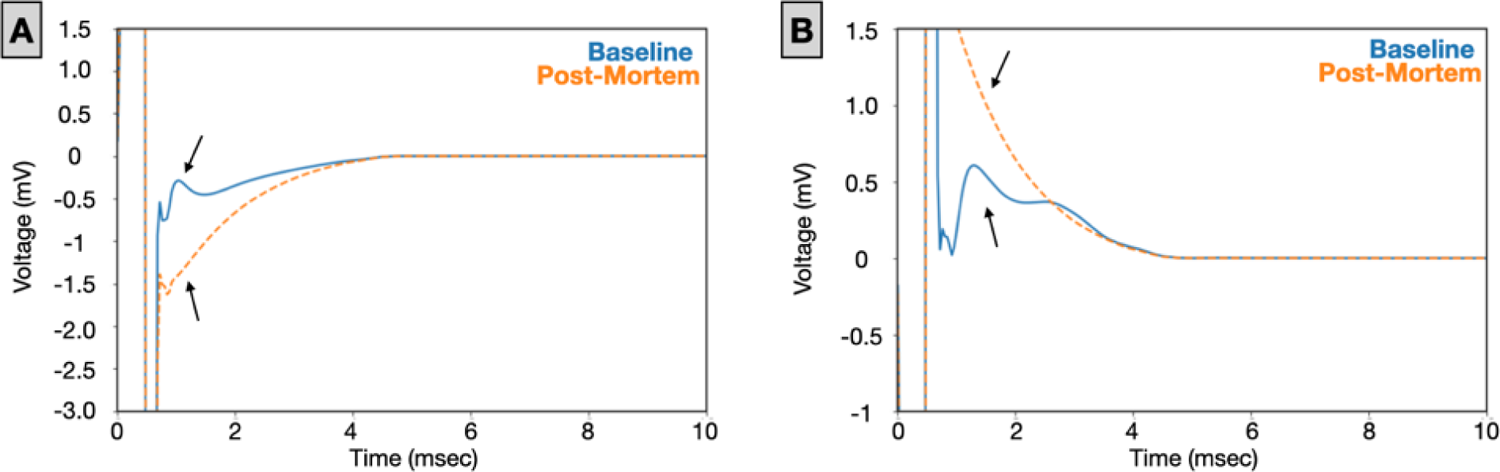
Representative, single contact ECAP recordings (contact 5) from a subject (subject 5) during baseline recordings (blue), as compared to post-mortem (orange). Stimulation cathode (10 mA) was swapped between contact 7 **(A)** and contact 8 **(B)**. Black arrows indicate the neural signal (baseline) or lack of neural signal (post-mortem). Note there is an A-beta neural component that appears to be ‘riding’ on the capacitive artifact caused by stimulation. The veracity of this apparent A-beta component can be confirmed by looking at the most distant recording contacts form the stimulation pair, where there is a clear separation of the neural component from the capacitive component of the stimulus artifact. However, it is problematic to assess what is ‘neural’ versus ‘artifact’ in situations where they are superimposed, which is common for most of the recording sites across the array.

In animals positively confirmed to have full elimination of stimulation evoked neural activity upon death, the capacitive artifact had a clear dependency on both distance from the stimulating electrodes to the recording pair and stimulation amplitude (Figure 5, Supplemental Figure 5 for all subjects). This artifact could consist of a post-stimulus positive deflection and then exponential decay which could last as long as 4-6 ms after the stimulus at the closest recording contacts and highest amplitudes tested (Figure 6). The shape of the capacitive artifact also changed with distance and amplitude, and at certain distances/amplitude it could easily be conflated with the P1/N1/P2 A-beta ECAP positive and negative deflections often used for closed-loop control (Chakravarthy et al., 2020; Parker et al., 2020). Capacitive decay at the electrode tissue interface is dependent on stimulation parameters and recording electrode coupling with the reference electrode. The post-mortem subject recordings followed a dose response curve typical of an N1-P1 peak response (Figure 6).

**Figure 5.**
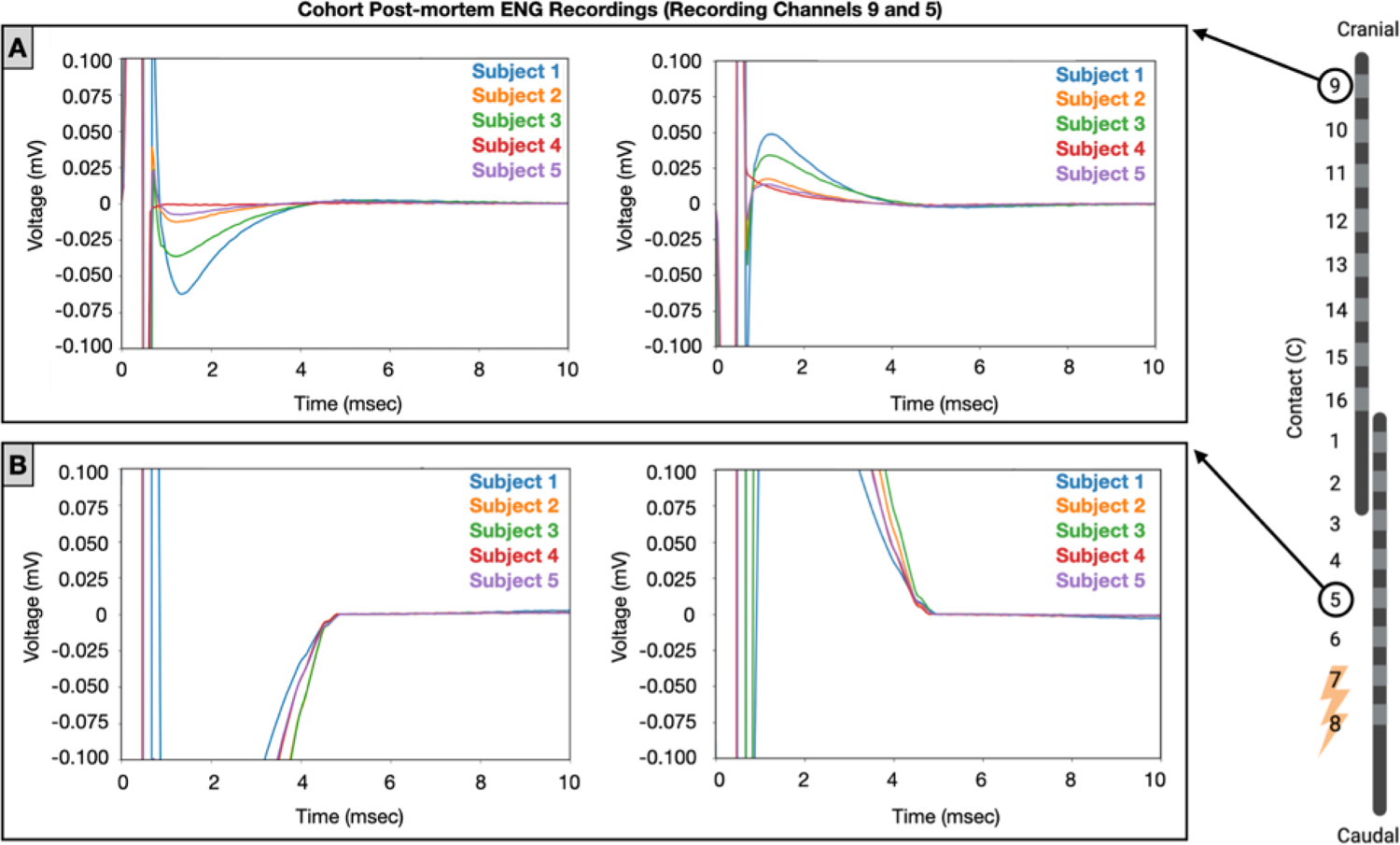
Post-mortem ECAP, single contact recordings for the cohort **(A)** away from (recording contact 9) and **(B)** near (recording contact 5) stimulating contacts 7 and 8 (electrode schematic). Stimulation amplitude was 10 mA for four of the five subjects as subject 1 had a lower maximum amplitude (7mA) due to muscle contraction. Stimulation cathode was swapped between contact 7 (left column A and B) and contact 8 (right column A and B). One subject did not have post-mortem stimulation recordings, and therefore is not included in this analysis. Note that at contacts closest to the stimulating electrode the capacitive artifact dominates the signal, however with more distance (contact 5) the signal starts to resemble neural peaks.

**Figure 6.**
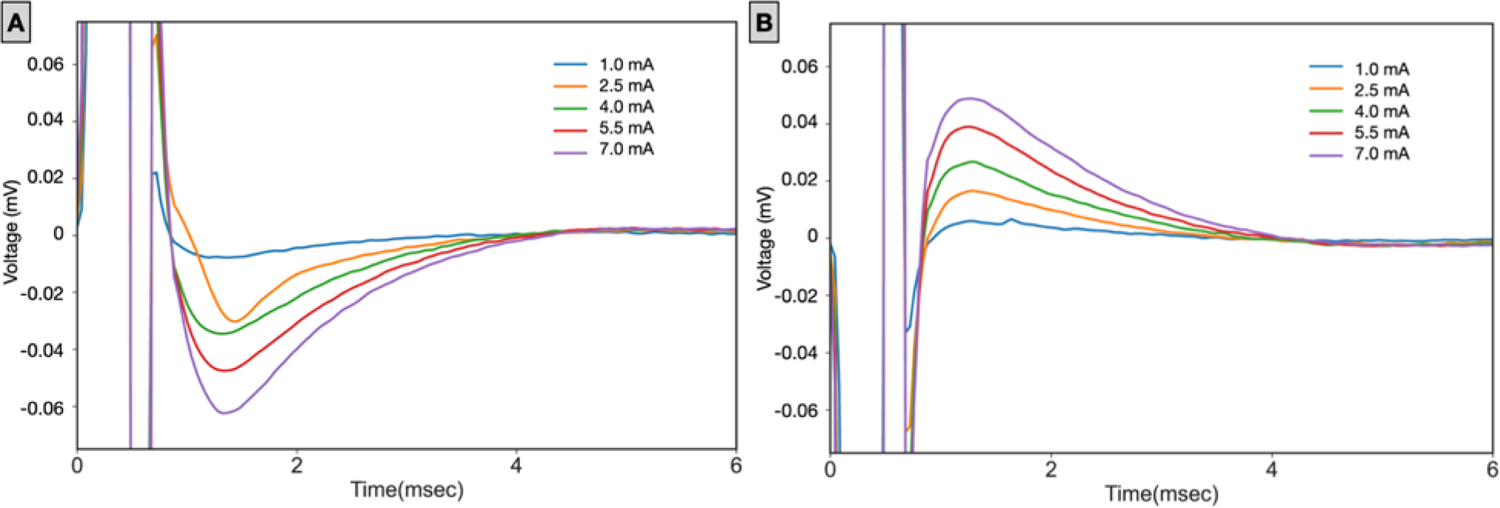
Representative (subject 1) post-mortem stimulation-induced capacitive artifact dose-response curve. **(A)** The dose response curve (recording at contact 9), when the cathode was at contact 7, and **(B)** contact 8 of the array. The P1/N1 created during post-mortem stimulations can resemble a P1/N1 in the dose-response curves of *in vivo* recordings, due to capacitive decay in the stimulation artifact (Supplemental Figure 5, all subjects DRCs).

### g. Dependency of Recorded ECAP Components on Respiration and Heart Rate

Visual inspection of the trace-by-trace recordings of evoked activity suggested a non-gaussian distribution of the recorded A-beta and EMG signal in most animals. Prior studies have shown that the amount of cerebrospinal fluid (CSF) at a spinal level between the dura and the underlying cord fluctuates as a function of inspiration/expiration and systole/diastole (Dreha-Kulaczewski et al., 2015; Friese et al., 2004). This could potentially change the coupling between the stimulating electrodes and the underlying cord, as well as between the fiber source of the ECAP and the recording electrodes. In addition, recording electrodes are very susceptible to motion artifact, in which the capacitive Helmholtz layer established at the electrode-tissue interface is disturbed by motion, creating an artifactual recording signal coincident with the source of motion (Giancoli, 1998; Ludwig et al., 2006, 2009; Michelson et al., 2018; E. Nicolai et al., 2018; Tam & Webster, 1977; Verma, Knudsen, et al., 2023; Wartzek et al., 2011). Consequently, we hypothesized that both the A-beta signal and EMG bleed-through recorded in the SCS ECAPs may vary cyclically as a function of motion due to breathing and heartbeat.

Figure 7 depicts representative ECAP data from one animal to illustrate the change of the ECAP component amplitudes over time during a pulse train. Figure 7A (wide view) and B (individual components) depict individual traces for each ECAP and EMG component over time compared to the mean average trace. Visual inspection of these individual traces is not consistent with a gaussian distribution around the mean trace but is more consistent with multiple separate groupings of traces depending on time. For example, note over the first 100 traces in Figure 7A most traces are above the mean for the positive peak of the second EMG component. Conversely, the later traces appear in the pulse train lower than the mean. During SCS at low frequencies, a decrease in amplitude of muscle responses over the course of a pulse train is often used as an indicator of activation of sensory afferents involved in a spinal reflex muscle response (Misiaszek, 2003; Palmieri et al., 2004).

**Figure 7.**
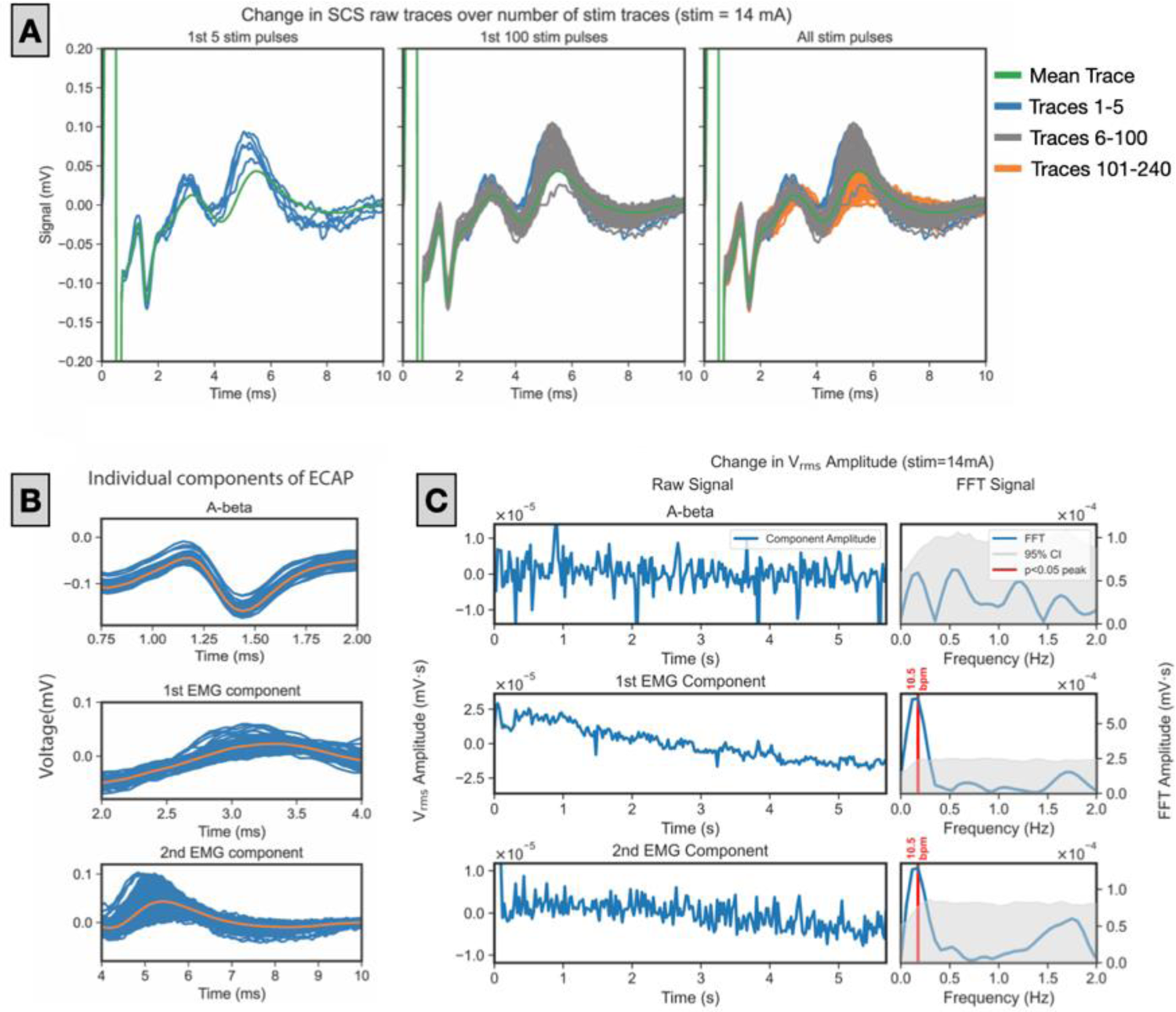
Representative evolution of ECAP components over the course of pulse train in one representative subject (subject 4). **(A)** green lines are the mean trace over all 240 pulses, the colored traces (blue, gray, and orange) indicate the raw traces with blue being the 1^st^-5^th^ stimulation pulses, gray being the 6^th^-100^th^ stimulation pulses, and orange being 101^st^-240^th^ stimulation pulses, and **(B)** all individual traces (blue) for each ECAP component over time (millisecond) as compared to the mean average trace (orange). **(C)** Change in the V_RMS_ over time for each ECAP, and EMG component, and FFT analysis for each.

A plot of the root-mean-square voltage (V_RMS_) amplitude of the A-beta and EMG responses over time during the pulse-train are also inconsistent with a random gaussian distribution, with clear oscillations in amplitude over time (Figure 7C). A Fast Fourier Transform (FFT) of the data in Figure 7D was used to identify the frequency components of the oscillatory activity. Clear peaks are evident in the 10-20 breaths per minute and 60-100 beats per minute range, consistent with the breathing rate and heart rate of the animal, respectively.

Figure 8 depicts V_RMS_ amplitude of each ECAP component and the associated FFT for every animal across the cohort. In every case, there is a large significant peak in the 10-20 breaths per minute range, consistent with the pre-selected breathing rate for the intubated animal. In addition, there is often a large secondary peak that corresponds to the approximate heart rate for the animal, but often not large enough to reach significance (See methods, Section D: Data Analysis for bootstrap technique used to determine FFT confidence interval for a gaussian distribution). It is notable that some of the V_RMS_ plots for the EMG components in some of the animals decrease over time during the pulse train (See Subject 5), consistent with the activation of a spinal reflex. However, in other cases the V_RMS_ increases over time (Subject 1) or has a complex appearance with an initial ‘decrease’ and then a general increase over the pulse train (Subject 7). There are additional smaller peaks evident in the FFT in many animals as well, which may be consistent with other sources of motion observed during the experiment such as tremor, which is common in anesthetized animals, or motion induced by contraction of large back muscles associated with stimulation.

**Figure 8.**
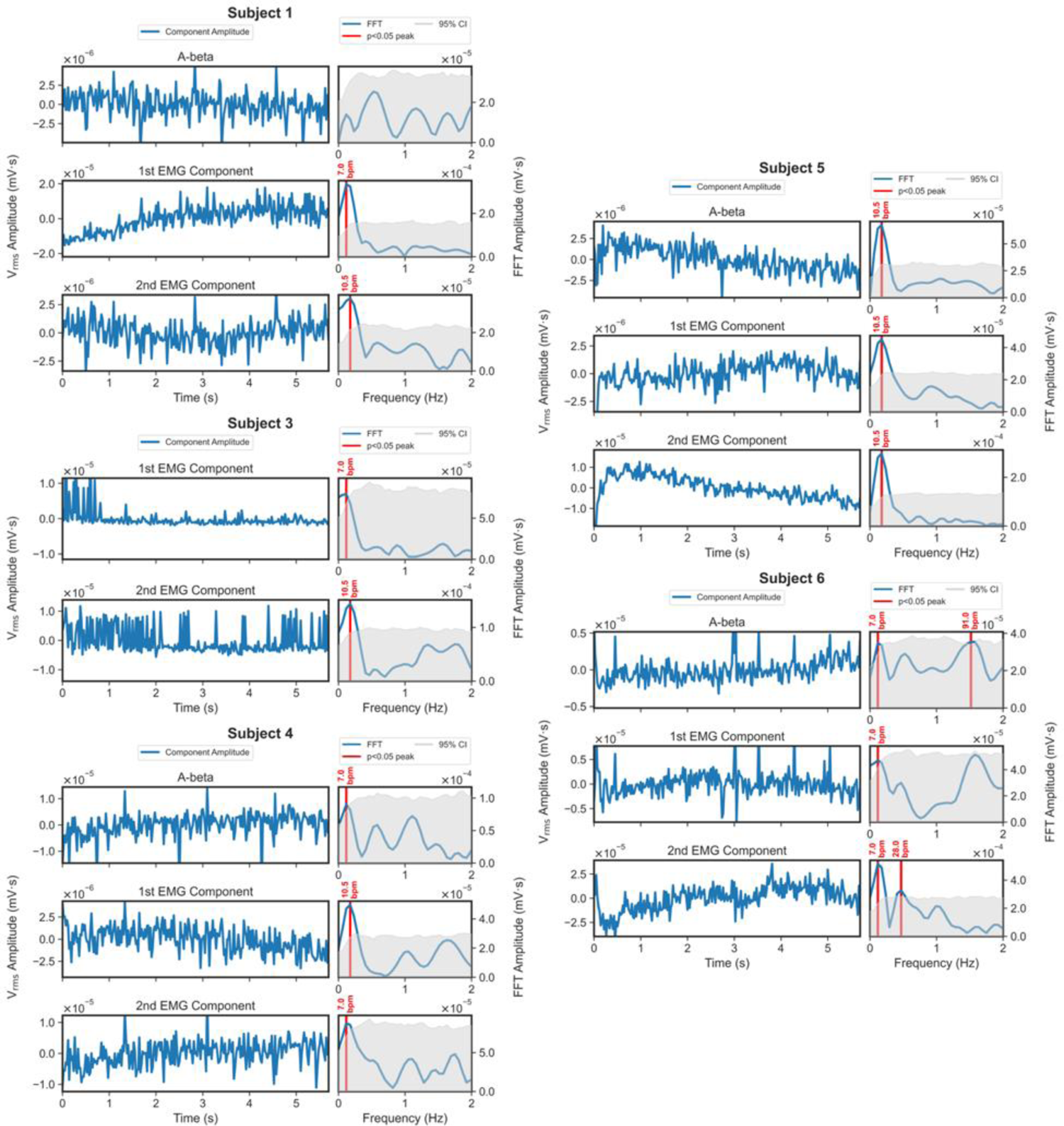
Examples of changes in V_RMS_ of SCS ECAP components across the cohort. Subject 2 was not included in this cohort analysis, as no EMG bleed-through was evident in that animal. Subject 3 A-beta component analysis was not included, as at even the most cranial recording locations the A-beta component was not sufficiently separate from the capacitive artifact to disambiguate it from the stimulus.

### h. Differential Dose-Response Curve for Early versus Late A-beta Response

After positively identifying the artifactual ECAP components, we noted a consistent secondary component in the A-beta range that appeared at the highest current amplitudes tested (Figure 9). The dose-response curve for these two peaks varied, wherein the first A-beta peak threshold was typically observed at 2-3 mA and the threshold for the secondary component was typically between 7-10 mA (Figure 10). Interestingly, this secondary component occurring only at higher amplitudes would often have a shorter latency than the first component evident at lower amplitudes. This result is surprising as the threshold for activation of a nerve fiber is inversely proportional to its diameter (larger diameter equals lower threshold), whereas the conduction velocity of the action potential propagated by the nerve fiber is proportional to its diameter (larger diameter equals faster conduction velocity), all other things being equal. These data would suggest that at higher current amplitudes there is a secondary point of activation closer to the recording electrodes. This could potentially be due to activation starting to occur at the anode, recruitment of an additional population of fibers based on anatomy, or engagement of a spinal reflex and associated delay (Gmel et al., 2023). For example, at lower thresholds activation may occur at the more superficial rootlets closer to the epidural electrodes due to shunting of CSF, and at higher thresholds of activation of the dorsal column may also occur as the action potential initiated at the rootlets would travel farther than an action potential initiated directly under the electrode.

**Figure 9.**
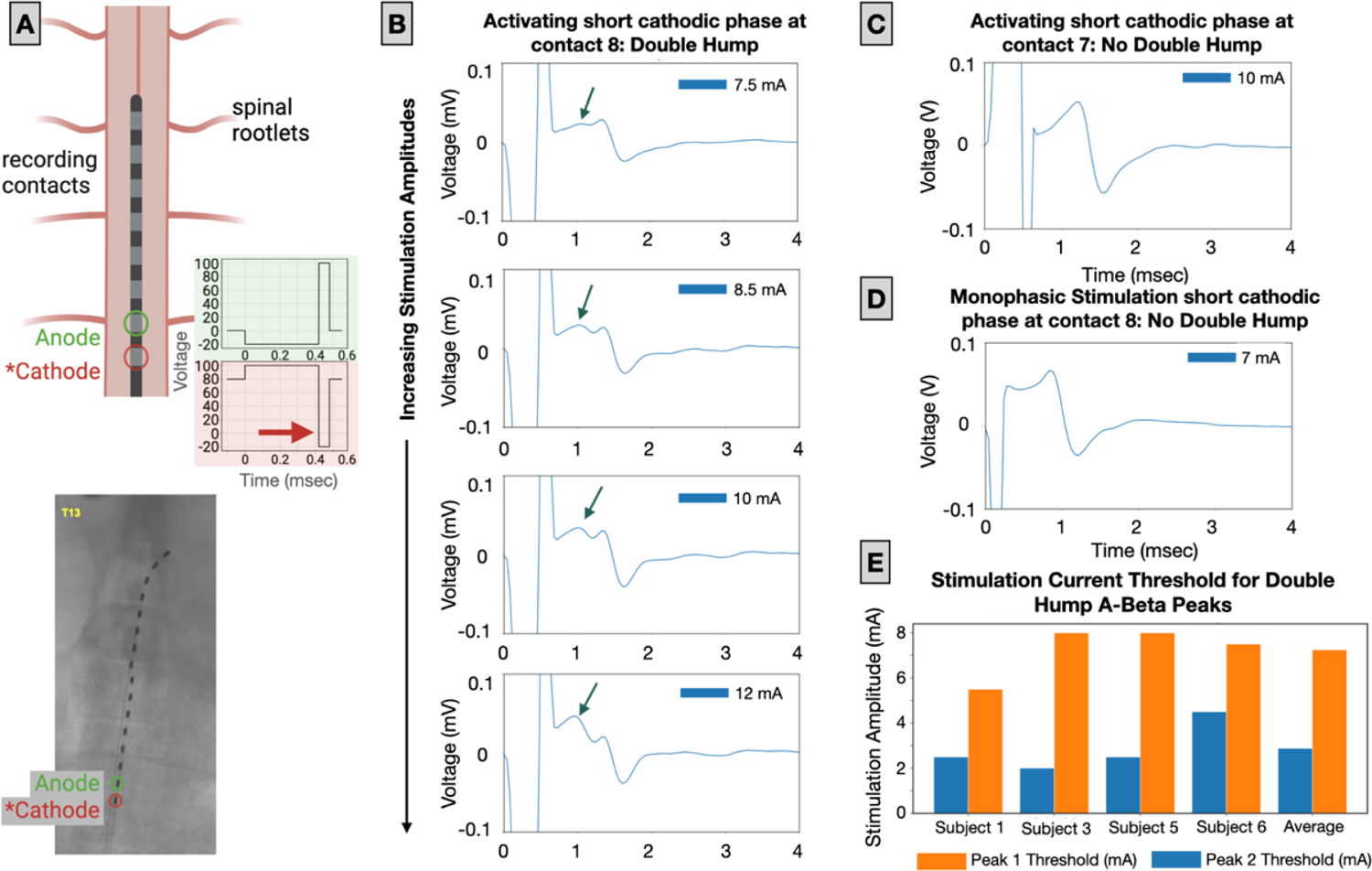
Evolution of the double hump in one subject (Subject 6) when the short cathodic pulse (*cathode) occurs at contact 8. All data shown was collected under the muscle paralytic (Vecuronium). **(A)** Schematic showing the spinal cord and spinal rootlets (not to scale) with the activating short cathodic pulse at contact 8 (red arrow), below is the corresponding CT showing placement of the electrode on the spinal cord. **(B)** With the activating short cathodic pulse at contact 8, the double hump is present, with the shorter latency peak increasing as the stimulating amplitude increases. The dark green arrows indicate the short latency component, resulting in the double hump effect. **(C)** When the stimulating configuration is swapped (with the short cathodic pulse occurring at contact 7, there is not a double hump present, even at increased amplitudes. **(D)** When a short cathodic monophasic waveform is applied to contact 8 there is not a double hump present. **(E)** Across all subjects the activation threshold for the double hump is shown for both peak 1 and peak 2, as well as the average. The activation threshold of 2^nd^ peak was consistently higher than the 1^st^ peak in all animals.

**Figure 10.**
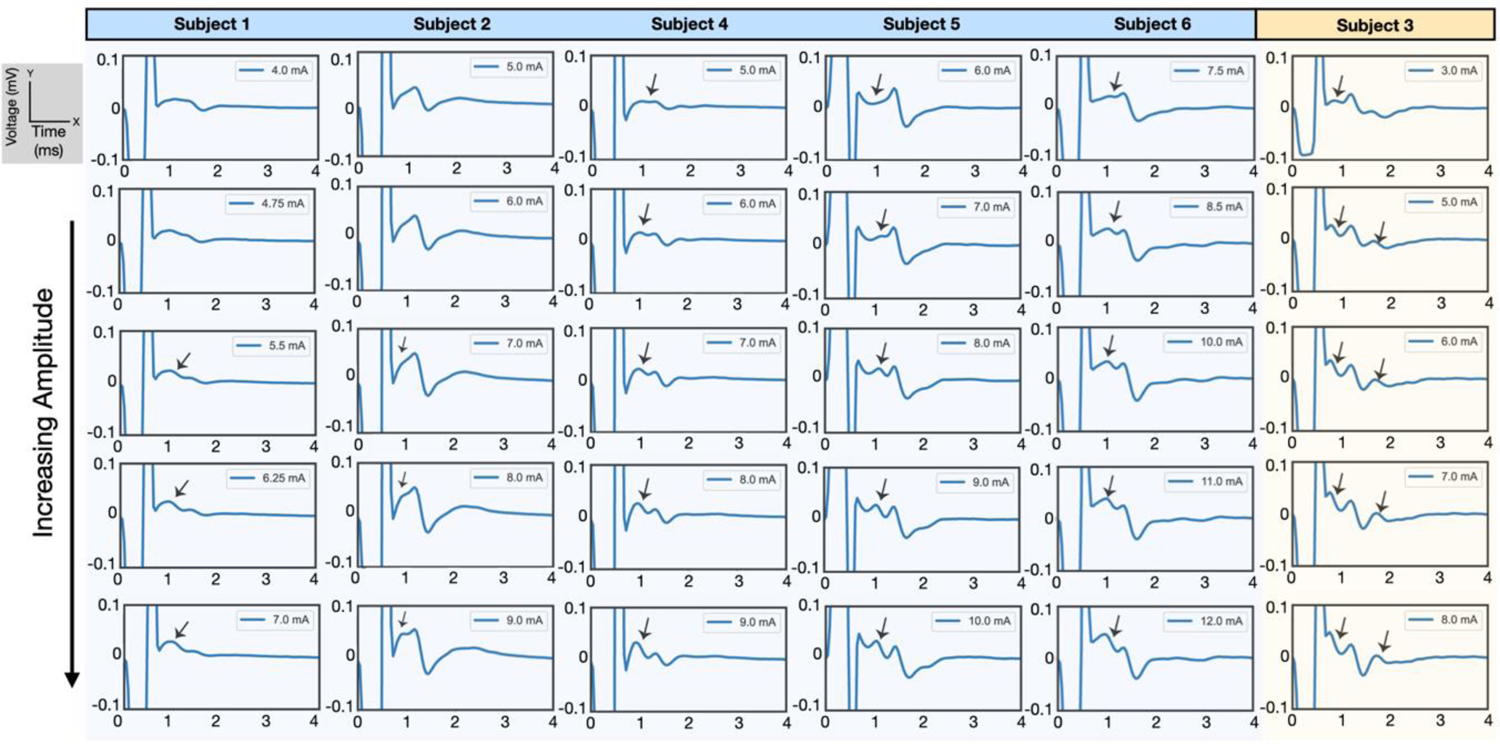
Evolution of double hump with increasing amplitudes across all animals (top to bottom). The second component was present in one stimulation configuration, short cathodic activating pulse on contact 8. The second shorter latency component appeared only as amplitude was increased in 5 of 6 subjects (blue). In one animal (yellow, subject 3), even at the lowest amplitudes given, all three neural components were present, so we were unable to determine the order of appearance as a function of current dose. The second component was verified to be neural by carrying out the analysis under vecuronium. [Supplemental Figures 6 & 7]

To preliminarily assess these two possibilities, we compared ECAPs recorded following the short cathodic activating pulse at contact 7, and then at 8. When the short cathodic phase was placed further from the recording electrodes (contact 8) we observed the ‘double hump component’ in the A-beta recordings at the highest current amplitudes (Figure 9). When the short cathodic phase was placed closer to the recording electrodes (contact 7), no ‘double hump’ was observed. These data are more consistent with the secondary A-beta component arising from concurrent activation at the anode at higher thresholds; however, this should be explored more deeply in future studies. The presence of the double hump was noted to occur at higher stimulation amplitudes across all animals (Figure 10). The second component was only present in one stimulation configuration, (short cathodic phase further from the recording contacts). In five subjects, a longer latency hump in the A-beta conduction velocity range was evident at lower currents before a shorter latency component eventually appeared. In one subject, three components/humps in the A-beta range were evident at even the lowest dose applied, so we were unable to determine which component had the lowest threshold for activation. The second component was verified to be neural by carrying out the analysis under Vecuronium (Figure 9 and 10).

## 4. Discussion

The ECAP recordings obtained during ESRs following epidural SCS can be contaminated by 1) capacitive artifact, 2) EMG of surrounding muscles of the back, and 3) motion. In a swine model of epidural SCS we demonstrate the potential confounds, overlap of signals and changes to recordings as a result of motion (breathing, heart rate etc.). Proper controls, such as administration of a muscle paralytic, should be implemented to differentiate neural signal from sources of artifact during acute epidural SCS recordings.

Ideally, the A-beta signal in recorded ECAPs could be determined by calculating the expected latency using the conduction velocity of the fiber type and the known distance of the stimulating electrode from the recording electrode (Blanz et al., 2022; Erlanger, 1937; Manzano et al., 2008; E. N. Nicolai et al., 2020). Unfortunately, although the distance from the stimulating electrode and its paired return are known, the point of activation in time and space is an unknown. It is unknown when fibers are activated during the cathodic pulse, or depending on dose, if there is concurrent activation at the anodic return due to the formation of a virtual cathode. At increasing current amplitudes, the point of activation along a fiber can move progressively further from the electrode as well (Mortimer & Bhadra, n.d.; Ranck, 1975). Changes in the site of action potential initiation, such as from the dorsal column or the rootlets as they enter the dorsal column, can change the distance an action potential travels from the point of initiation to the point of recording.

Given that the site of action potential initiation is unknown, it is common to use the propagation of the P1 or N1 component of the putative A-beta response from site to site across a recording array to estimate the conduction velocity. Our data demonstrate that there are several confounds that need to be considered when using just the propagation of a P1 or N1 component across the array to isolate neural signal from artifact. In our data set, we used two eight-contact electrode arrays placed end to end, creating a distance of approximately 105 mm from the center of the stimulating electrode pair to the furthest recording electrode, assuming a distance of 7 mm center to center between the arrays. Without an artifact rejection applied, A-beta components are frequently not separate and distinct from the stimulation artifact capacitive component until reaching the recording sites on the second contiguous

SCS electrode array, within the range of current amplitudes typically applied clinically, given an 80-µs cathodic phase (Figure 3). Moreover, components of the A-beta response can ‘disappear completely’ into the stimulation artifact at recording contacts two to four electrode sites away from the stimulating pair. As the distance between the stimulation and recording sites decreases and the ohmic stimulation artifact and its capacitive decay become larger, this can give the appearance of stereotypical P1 or N1 peaks that change their latency systematically at progressively distant recording locations.

Similarly, we demonstrate that a very short latency evoked EMG bleed-through in the 2 to 4 ms range – and therefore from a muscle group very close to the SCS stimulating contacts – can also be superimposed on the A-beta signal. As the two electrode arrays can span the local dipole source created by the muscle activation, the EMG representation via bleed-through in the SCS recordings increases amplitude at locations closer to the dipole source and can even flip polarity between recording electrodes closer to the positive or negative pole of the dipole (Humphrey & Schmidt, 1990). These changes across the electrode arrays can give the appearance of the stereotypical ‘phase delay’ due to an ECAP propagating across the same array (Figure 3). The existence of evoked EMG in the 2 to 4 ms range, suggesting very local muscle activation, is consistent with activation of the long muscles of the back, which should be explored more thoroughly in future studies. Activation of the long muscles of the back such as the longissimus or multifidus at relatively low current amplitudes suggests a potential source of off-target activation to be avoided, or even a substrate for possible benefits of SCS. Activation of the multifidus muscle, for example, has recently been FDA approved for the treatment of chronic back pain (Gilligan et al., 2023a, 2023b).

### a. Best Practices for Identification of Artifacts and Reporting of Data

The sources of artifact for ECAP recordings are well understood for other neuromodulation targets where ECAPs are commonly used to understand the activation of nerve fiber types responsible for therapeutic effects to optimize electrode configurations and stimulation parameters (Kikuchi et al., 2005; E. Nicolai et al., 2018; Rosahl et al., 2000; Verma, Knudsen, et al., 2023; Yoo et al., 2013). The most reliable method to determine the neural origin is to transect the nerve pathway between the stimulation and recording sites (Kikuchi et al., 2005; E. Nicolai et al., 2018; Rosahl et al., 2000; Verma, Knudsen, et al., 2023; Yoo et al., 2013). However, this may be problematic for an SCS experiment as it would involve transection of the spinal cord itself. As an alternative in acute animal experiments, we recommend using the administration of a muscle paralytic with concurrent EMG recordings to confirm sufficient muscle block via the elimination of directly recorded EMG responses. Slow evoked responses (termed eSAPS), that are not-artifact and non-myogenic, have been described in a rodent model of SCS - and attributed to synaptic currents within the dorsal horn (Sharma et al., 2023). Physiological slow potentials from the spinal cord have been characterized for decades (Shimoji et al., 1982; Yates et al., 1982) and even informed the development of gate theory (Bikson & Sharma, 2024). However, our study shows that to the extent eSAPS can be recorded clinically, their identification will need to account for the artifacts described here. In addition, we also conduct recordings ∼5-10 minutes after euthanizing the animal to get a representation of stimulus artifact at different dosages resembling the live *in vivo* experiment as closely as possible. This approach also requires concurrent EMG recordings to ensure induced neural activity has ceased, as well as the elimination of A-beta components at the most distal recording sites.

Unfortunately, our recommended methods would not be effective for chronic animal experiments or human studies. In these cases, we still recommend calculating the nerve conduction velocity by propagation of the P1 or N1 peak of the ECAP across the recording array. Although we have demonstrated here that this is not completely infallible, putatively due to the activation of local sources of EMG or stimulation artifact, this is still a useful tool to help disambiguate sources of artifact from the neural signal. Ideally, the use of recording sites with sufficient distance from the stimulation sites allows for clear separation of the neural signal from the capacitive component of the stimulation artifact. This clearly separate neural signal can then be traced by its phase delay along the electrode array, moving closer to recording sites; this may help identify neural signal, as well as when the neural signal is no longer evident due to complete masking by the stimulation artifact.

Observing the propagation of neural signal across the recording SCS array at different current amplitudes and stimulation configurations can also be a useful tool to verify neural signal. This can help determine if the dipole source of the muscle causing EMG bleed-through is spanned by the recording array, which can in turn, lead to progressively decreasing amplitude of the EMG bleed-through or even flipping of polarity across the recording array. This can create the artifactual appearance of progressive phase delay across the recording array due to the ‘tip of the iceberg’ phenomenon. As more of the ‘tip of the iceberg’ is visible above the noise, the starting and ending point of an EMG peak (the edges) may appear to systematically shift. As suggested by Figure 9 and 10, at increasing doses it may be possible to observe a second A-beta component due to activation at the anode when the short anodic phase of stimulation is closer to the recording electrodes.

The examples provided in this study demonstrate the potential confounds of using a small temporal window of ECAPs at one recording site to assess the likelihood that evoked responses are of neural origin or caused by side effect/artifact. We would suggest that simultaneously recording average ECAPs and trace-by-trace plots across the recording array should be provided to accurately potential artifact contamination. These data should be provided at both the common ‘zoomed’ in temporal window (i.e. 0 - 2 ms post stimulation pulse) and at expanded windows out to at least 10 - 14 ms. These longer windows are necessary to identify multiple potential muscle sources of EMG bleed-through from the back and legs and, depending on frequency of stimulation, to assess if delayed EMG artifact may be contaminating a subsequent biphasic stimulation pulse response.

### b. Impact of Electrode Design, Instrumentation, and Post-Processing on Artifacts

The artifacts highlighted in this study are heavily dependent on many factors, including electrode design, stimulation instrumentation, stimulation waveform/configuration, recording instrumentation, referencing, post-instrumentation filtering, artifact post-processing, and several environmental factors. ***Therefore, the data presented in this paper are very specific to stimulating and recording from two epidural SCS electrode arrays placed end to end to maximize the distance from the stimulation and recording sites, when stimulating with an asymmetric biphasic pulse.*** Given the prevalence of these artifacts, it is common for other neuromodulation targets to use a variety of techniques to minimize their impact on the recorded ECAPs, including blanking the stimulation window, quick settle amplifiers (Elyahoodayan et al., 2019; Samiei & Hashemi, 2021), forward masking (Baudhuin et al., 2016; Kirby & Middlebrooks, 2010), exponential subtraction of the stimulus artifact (Chakravarthy et al., 2022), inverted polarity stimulation and subtraction, and tripolar recording techniques (Yoshida et al., 2009). Filtering can also have an effect on the capacitive component of the stimulation artifact. In post-mortem recordings without filtering the capacitive artifact can appear longer and contain ringing (Figure 11). Although these artifact reduction strategies can be very useful in minimizing the contamination of evoked neural components in ECAP recordings, their effectiveness can vary between stimulation amplitudes, configurations, and over time. Moreover, these techniques which rely on ‘subtracting’ out the stimulation artifact from the recorded neural signal when applied to stimulation artifact without EMG/ECAPs can leave a residual component that may also mimic the appearance of neural signal. Therefore, regardless of the strategy used to minimize the impact of the stimulation artifact and EMG, it is still important to take appropriate steps to confirm the sufficiency of these methods. Wherever possible this should include transections, application of muscle paralytics, and inclusion of complete data sets sufficient for the reader to make their own judgment on the extent to which artifacts may be contaminating the neural signal^2^.

**Figure 11.**
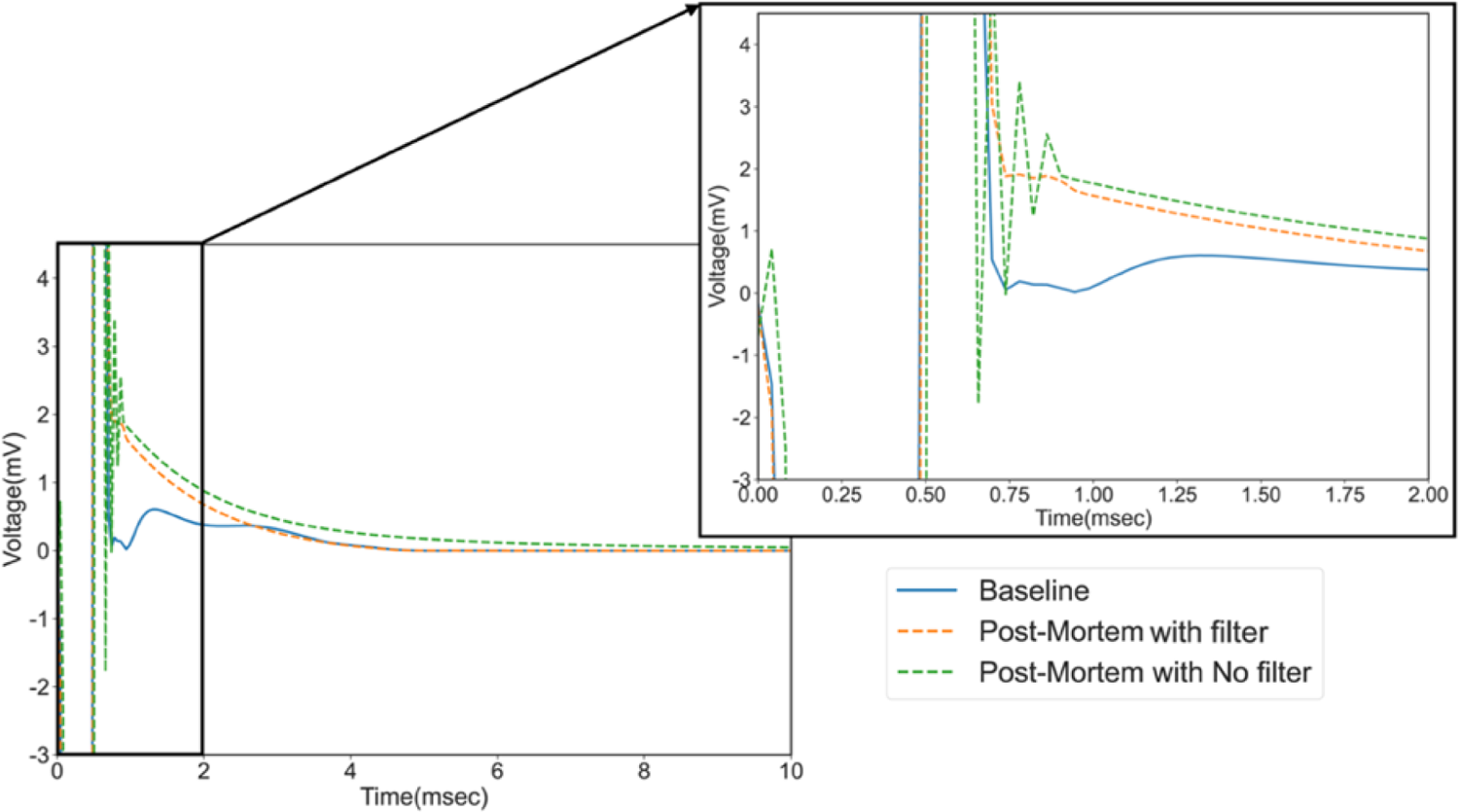
Impact of filtering on stimulation (10 mA) artifact capacitive decay. Baseline and post-mortem recording from a representative subject (subject 4), with zoomed window showing longer capacitive artifact and filter ringing (inset). Cathode is on contact 8. As can be seen in the dotted green trace the intrinsic hardware filters cause some filter ringing. When a post-hoc filter is applied this ringing is mostly removed but can leave the appearance of a small bump (dashed orange) that could be mistaken for an a-beta.

### c. Intermittent Sources of Artifact

Even during an acute experimental paradigm in a well-controlled setting with an anesthetized animal, occasional transient artifacts can be observed across the recording array. This may be due to 1) powered equipment from within or near the surgical room turning on/off causing transient sources of radiofrequency (RF) noise, 2) ground loops created when recording both SCS ECAPs and multiple concurrent EMG pairs, and/or 3) intermittent tremor which is common in an anesthetized pig. The appearance of these transient artifacts can be difficult to predict, as they depend on both the instrumentation being used and any post-hoc filtering performed. A step function input into a filter such as those caused by transient motion or nearby sources of RF can cause the appearance of a gradually decreasing oscillatory signal at the lowest frequency of the high-pass cut-off, commonly referred to as filter ringing (Heffer & Fallon, 2008; E. Nicolai et al., 2018; Verma, Knudsen, et al., 2023). Consequently, filter ringing can create the false appearance of ECAP components. Although care was taken in this study to use post-hoc filtering techniques known to be less susceptible to the ringing effect (see Methods) (Bovik, 2005), some small ringing was observed due to the intrinsic hardware filters of the TDT system used for recording and after large transient artifacts (Supplemental Figure 8). Additionally, transient artifacts can be mitigated using signal averaging. Standard clinical applications that use pulse by pulse responses for closed loop applications may not take advantage of this averaging effect, and therefore may result in an incorrect adjustment to stimulation due to these transient artifacts.

### d. Implications of Motion Related Oscillations in ECAP Components

There are multiple potential sources for the oscillations in V_RMS_ amplitude observed in the recorded A-beta signals and EMG bleed-through in this study (Figure 7 and 8). Prior studies have shown that the amount of CSF between the epidural surface and the underlying cord cyclically changes as a function of both respiration and heartbeat (Delaidelli & Moiraghi, 2017; Dreha-Kulaczewski et al., 2015; Friese et al., 2004; Lloyd et al., 2020). This, in turn, changes the distance of the epidurally placed stimulating electrodes to the underlying nerve fibers potentially causing cyclical differences in activation, and also changes the distance of the epidural recording electrodes from the neural sources of evoked activity. Moreover, respiration causes the chest wall to expand and contract, bringing potential muscle sources of EMG bleed-through cyclically closer to and further from the epidural SCS recording array. A common misconception is that these motion related artifacts can be eliminated by filtering selections because they occur periodically at low frequencies. Unfortunately, as the artifact is created by disturbance of the Helmholtz layer between the electrode and underlying tissue and subsequent reestablishment of the Helmholtz layer, these motion artifacts have higher frequency components which distort ECAP components in the bandpass range (Bard et al., 1980; Giancoli, 1998; Merrill et al., 2005; Michelson et al., 2018; E. N. Nicolai et al., 2020; Tam & Webster, 1977; Wartzek et al., 2011).

Our data suggest that the EMG bleed-through across the cohort was the most susceptible to these cyclical oscillations, with the largest power in the frequency band matching the ventilation rate maintained across the cohort. This would suggest that expansion/contraction of the chest wall during breathing causing muscles to move progressively closer to and further from the SCS recording array was the most common source of observed oscillations in EMG bleed-through. However, we periodically found similar cyclical oscillations in the recorded A-beta amplitude of an unknown origin. Putatively, these may be caused by relative motion of the stimulation and recording electrodes to the underlying spinal nerve fibers, or the general motion of the SCS electrode array due to cardioballistic/respiratory motion. Similar motion artifacts linked to heartbeat and respiration have been observed in microneurography recordings from peripheral/cranial nerves in a prior study, which were verified as motion artifacts by their continued presence even after nerve transection between the stimulation and recording electrodes(Verma, Knudsen, et al., 2023). It should also be noted that the voltage signal created by electrical activity during heartbeat, commonly recorded in electrocardiogram (EKG), is of sufficient magnitude to be evident in ECAP recordings, if insufficiently addressed by common mode rejection and filtering. The EKG voltage waveform has different frequency content then motion artifact created by the motion of the heart and CSF changes. These changes are linked to the cardiac cycle disturbing the Helmholtz layer and/or stimulation/recording distances to neural tissue. As a result, the frequency components of these cardioballistic artifacts are not explicitly addressed by filtering designed to minimize EKG voltage bleed-through in the ECAPs.

It is likely that there is some small effect on the relative distance of spinal nerve fibers to the epidural stimulation/recording during the cardiac and respiratory cycles (Delaidelli & Moiraghi, 2017; Dreha-Kulaczewski et al., 2015; Friese et al., 2004; Lloyd et al., 2020). Considering that constant activation of A-beta fibers may help provide consistent pain relief, these changes in stimulation and recording coupling based on CSF changes during the respiration cycle and may need to be accounted for to optimize activation of the same population of A-beta fibers in a closed-loop paradigm.

### e. Implications for Recording and Interpretation of ECAPs for Closed-Loop Control in Real World Setting Outside of the Laboratory

Here we’ve shown that ECAP recordings are difficult to disambiguate from artifact in a very controlled experimental setting, whereby sources of artifact such as motion, and external equipment can be systematically identified, isolated, and mitigated. The use of ECAPs outside of controlled laboratory environments, as may occur during awake-behaving settings in the home, further complicates the disambiguation of recorded neural signals from multiple sources of artifact, especially during alterations in posture, cough etc., when the spinal cord is known to move relative to the epidural surface. This is especially problematic when these ECAP recordings are specifically used to detect and adjust for changes in evoked A-beta amplitude in a closed-loop system. The Medtronic Restore system previously used an accelerometer within the implantable pulse generator in an attempt to adjust stimulation during postural changes, cough, etc. (Kumar et al., 2014). Interestingly, even if the amplitude of the evoked A-beta fiber response is largely contaminated by motion artifact during these conditions in a real-world setting, the presence of this motion artifact itself could be a reliable predictor of motion at the SCS electrode array relative to tissue. Consequently, even an SCS ECAP recording that can be easily contaminated by motion artifact may be useful in terms of closed-loop stimulation paradigms to adjust for periods of relative motion.

It is important to note that dose-response curves of EMG bleed-through caused by an external muscle dipole source will be less sensitive than direct measures of EMG at the muscle. Depending on the distance and orientation of the dipole source, EMG bleed-through may only be evident at current amplitudes much higher than those needed to observe evoked EMG using direct recordings at the muscle. Therefore, it is important not to use EMG bleed-through measured during SCS ECAP recordings as a stand-alone proxy for unwanted off-target activation for a given stimulation electrode configuration and set of stimulation parameters.

### f. Limitations

Several limitations need to be considered when interpreting these results. The first limitation is the effects of anesthesia on the subjects during ESRs. Subjects in this study were intubated and maintained on isoflurane gas. Regardless of its safety, isoflurane can be a depressant on the cardiovascular system, and result in decreases in blood pressure (Kohn, 1997). Additionally, it has been shown that isoflurane can dampen EMG responses to SCS, as compared to alternative injectable anesthetics (Toossi et al., 2019). This highlights the necessity for awake, chronic studies to determine the appropriate stimulation parameters.

Second, the pig model has been deemed one of the more applicable research models when comparing nerve diameters, and fascicular organization to that of humans (Settell et al., 2019). Regarding spine and spinal cord similarities, swine are also similar to human in the shape and organization of the vertebral canal. Though human vertebral canals have a larger width, the overall patterns for sizing between different vertebral levels are similar (Busscher et al., 2010; Dath et al., 2007). This slight difference in anatomy may affect direct translation of stimulation parameters and should always be considered when translating results.

Third, the placement of leads was conducted during a laminectomy procedure. Therefore, electrodes were visible, which allowed for direct placement at the midline. In the hospital setting, leads are generally placed percutaneously. Therefore, leads may not be in a location that is directly comparable to an animal model. A reference was placed in adipose tissue near the spinal cord to avoid differential subtraction of the ECAPs that occurs when both the working and reference electrode are on the same epidural array (Verma et al., 2023). However, using a reference as close as possible to the working electrode will increase the similarity of stimulation artifact and EMG bleed-through, and improve subsequent common mode rejection (Verma, Romanauski, et al., 2023). This may have exacerbated the appearance of EMG bleed-through and stimulation artifact, but based on previous studies by our group this effect is relatively minimal (Verma, Romanauski, et al., 2023).

Importantly, a charged balanced asymmetric waveform with the long anodic phase preceding the short cathodic phase was used in this study instead of a charged balance cathodic leading pulse, used clinically. This was chosen to limit the period of stimulation resistive and capacitive artifact contaminating the ECAPS immediately following the cathodic pulse, presumed to activate the nervous tissue. Anodic leading asymmetric pulses have been proposed to inhibit/inactivate large diameter fibers and/or fibers close to the electrode prior to a cathodic pulse in certain situations (Grill & Mortimer, 1995; McIntyre & Grill, 1999, 2000). This could have raised the threshold for cathodic activation in our study, leading to larger capacitive artifacts. However, the range of amplitudes tested and demonstrated to show a clear A-beta response were within the normal clinical ranges given an 80 µs cathodic pulse (Yearwood et al., 2010).

Finally, following the placement of electrodes, patients are allowed a healing-in period, where postural changes and scarring may result in movement of the electrodes. Clinical trial data on spinal cord stimulation shows that when a patient bends, sleeps on their back, or performs other general day-to-day movements, it can cause changes in activation threshold for dorsal column and dorsal roots (Mekhail et al., 2020). We are planning future chronic pig studies to explore the effects of postural changes (e.g. lying supine vs. prone) on the activation threshold for back muscles and the dose-response curves and recordings of ESR.

## 5. Conclusions

In conclusion, many closed-loop paradigms rely on ESRs to monitor the activation of the dorsal column fibers, while attempting to optimize their activation and optimizing their stimulation paradigm to maximize the therapeutic window. However, recordings from the spinal cord are subject to a multitude of artifacts and contamination, making them a moving target for closed-loop SCS. Here we positively confirmed the sources of these artifacts in a swine model of SCS as being, 1) stimulation artifact interfering with A-beta fiber recordings near the electrode, 2) artifact from muscle activation (EMG), and 3) respiration/cardiac cycle. Each of these components exerts some effect on the SCS recording, with dependency on the relative recording location from the stimulating location. All these components need to be considered when using patient recording data for closed-loop machine learning training algorithms. Even using modern tools like large datasets and generative artificial intelligence (AI) can lead to erroneous conclusions if proper controls are not considered when understanding the underlying complications in datasets.

Future studies should consider the effect of stimulation artifact and motion, including heart rate, respiration, and movement, on the recording and associated fiber signals. Additionally, proper controls, such as muscle paralytics, should be utilized to differentiate neural signal from EMG artifact in recordings. By appropriately addressing each of the illustrated confounds to spinal cord ECAP recordings, these signals can hopefully be more effectively used to improve both mechanistic understanding and therapeutic outcomes from closed-loop devices.

## Acknowledgements/Funding/Disclosures

The authors would like to thank Sarvesh Periyasamy for his assistance during biplane CT imaging and fluoroscopy, and Dr. Margo Straka and Dr. Marom Bikson for their helpful suggestions and feedback.

This study was funded by Abbott Neuromodulation. JKT is a paid consultant for Presidio Medical, Inc. NV was an employee of Abbott Neuromodulation and BioCircuit Technologies during the completion of this work. NV is a stockholder in NeuronOff and NeuraWorx. NV is currently an employee of Presidio Medical. KAL is a co-founder and equity holder for Neuronoff, Inc. KAL is also a co-founder and equity holder of NeuraWorx. KAL is a scientific board member and has stock interests in NeuroOne Medical Inc. KAL is also a paid member of the scientific advisory board of Abbott and Presidio Medical, and a paid consultant for the Alfred Mann Foundation, ONWARD and Restora Medical. KAL and AJS are co-founders of Neuronoff Inc. AJS is a consultant to Neuronoff Inc. SFL receives research support from Abbott Neuromodulation, Medtronic plc, Neuromodulation Specialists LLC, and Presidio Medical Inc.; is a shareholder in CereGate, Hologram Consultants LLC, and Presidio Medical Inc., and is a member of the scientific advisory boards for Abbott Neuromodulation, CereGate, and Presidio Medical Inc. IL receives research support from Abbott Neuromodulation; DJW is a co-founder and shareholder of Reach Neuro, Inc. and Bionic Power, Inc. D.J. Weber has stock interests in NeuroOne Medical, Inc., Neuronoff, Inc. D.J. Weber receives research support from Meta, Inc. ERE, MZ, and HP were all employed by Abbott neuromodulation during the completion of this work. The remaining authors declare that the research was conducted in the absence of any additional commercial or financial relationships that could be construed as a potential conflict of interest.

## Footnotes

1 Direct activation occurs when the electric field applied epidurally activates the ventral efferent roots/rootlets closest to the electrode, in part due to significant current shunting of the cerebral spinal fluid in large animal. Indirect activation of ventral motor efferents occurs when sensory afferents are directly activated by the applied electric field that participate in a reflex arc.

2 IE only showing ‘representative’ data from a single contact and/or very short window after the stimulation artifact, instead of including full data sets in the supplement.

