## Supplemental Materials for "Epidural Spinal Cord Recordings (ESRs): Sources of Artifact in Stimulation Evoked Compound Action Potentials"

### Supplemental Figures

**Supplemental Figure 1**: Microdissection of the swine (subject 2) demonstrating that T15/L1 spinal nerves exit and innervate the long muscle of the back, as well as the intercostal muscle.

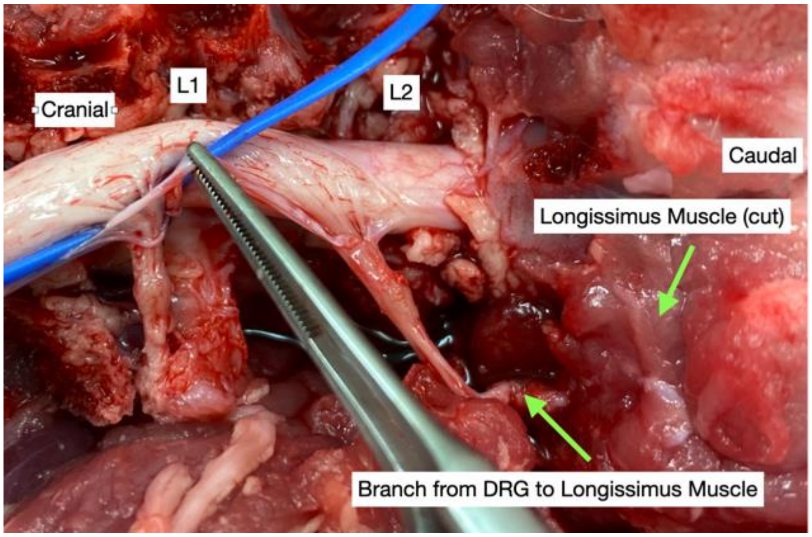


**Supplemental Figure 2:** Bimodal distribution of outlier traces in one subject. Pulse-by-pulse and averaged ECAP recordings following 9 mA of stimulation with cathode at contact 7 **(A)** and cathode at contact 8 **(B)**.

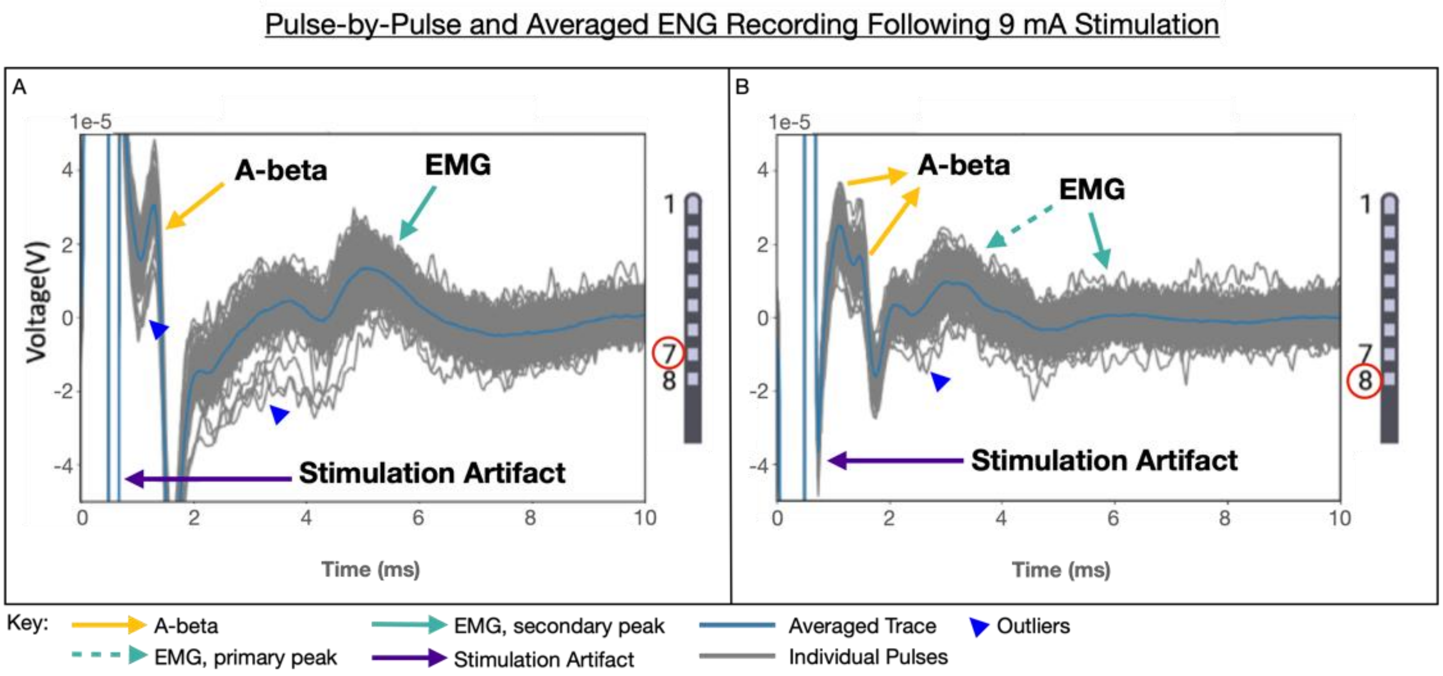

**<u>Supplemental Figure 3</u>:** Cohort Baseline vs Vecuronium ECAPs after 9-10mA Stimulation (All Animals). Orange indicates the baseline recording, and blue dashed line indicates the recording under vecuronium.

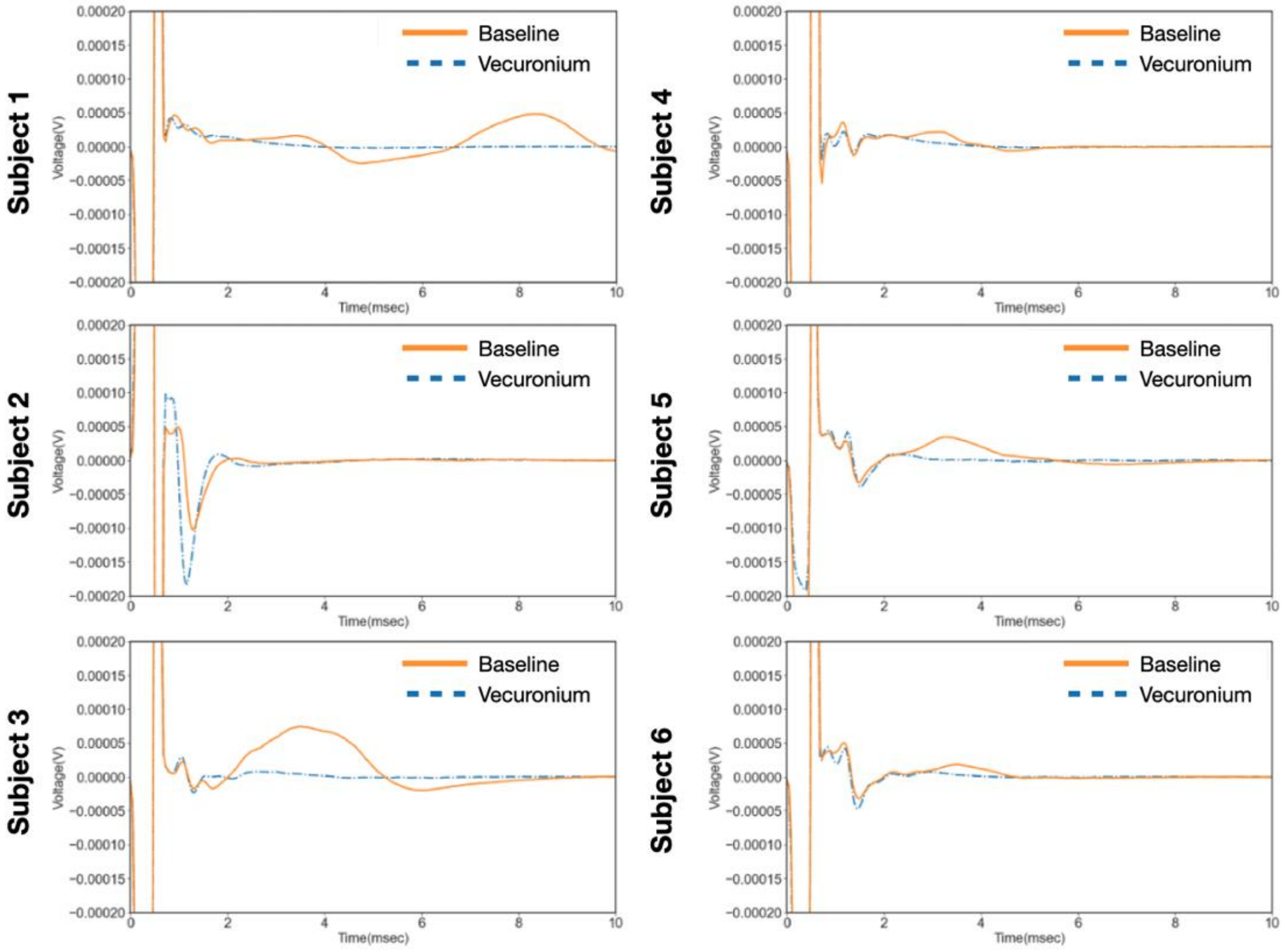

**Supplementary Figure 4:** Individual animal ECAP recordings across the array for both stimulation configurations (cathode at 7 vs. 8) at baseline and following vecuronium. Amplitudes are as noted in individual plots.

**Animal 1 (cathode on 7):** Baseline - Recording Channels 1-6

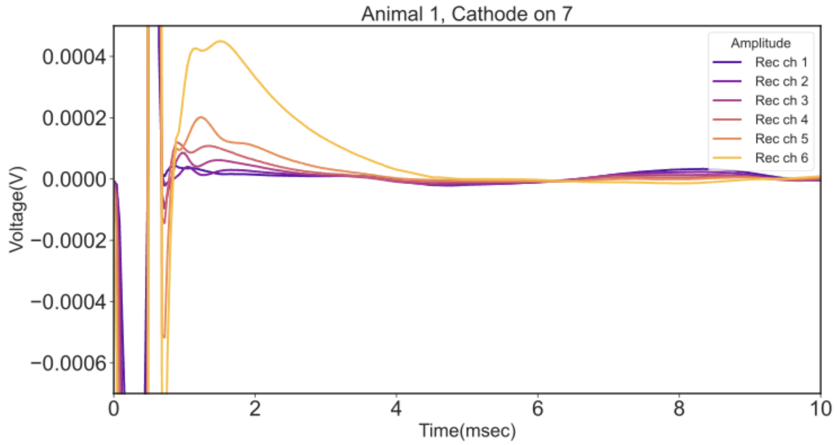


**Animal 1 (cathode on 7):** Vecuronium - Recording Channels 1-6

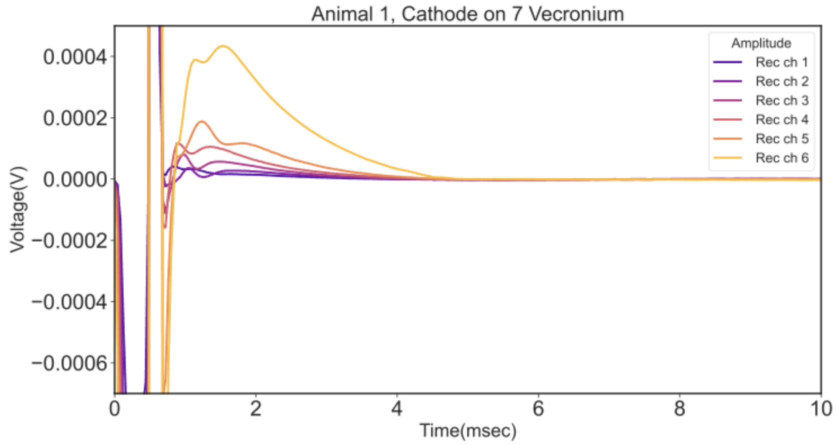


**Animal 1 (cathode on 7):** Baseline - Recording Channels 9-16

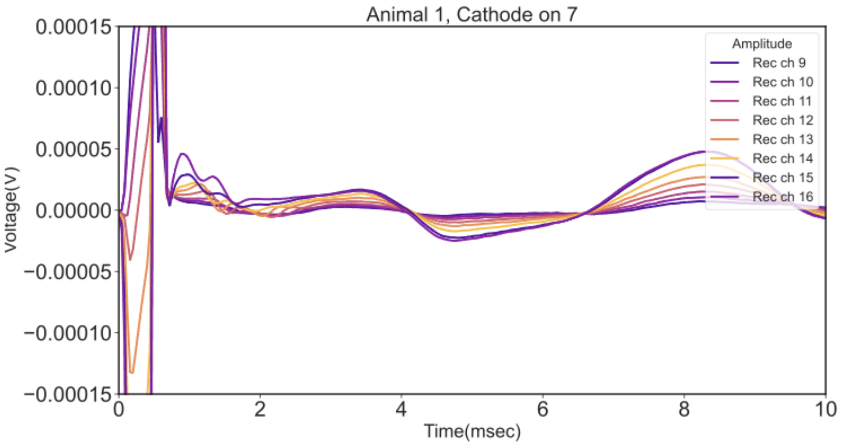


**Animal 1 (cathode on 7):** Vecuronium - Recording Channels 9-16

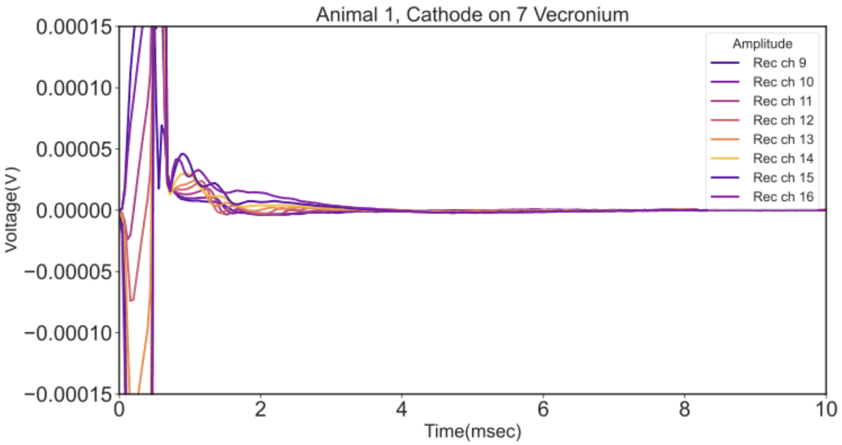

**Animal 1 (cathode on 8):** Baseline - Recording Channels 1-6

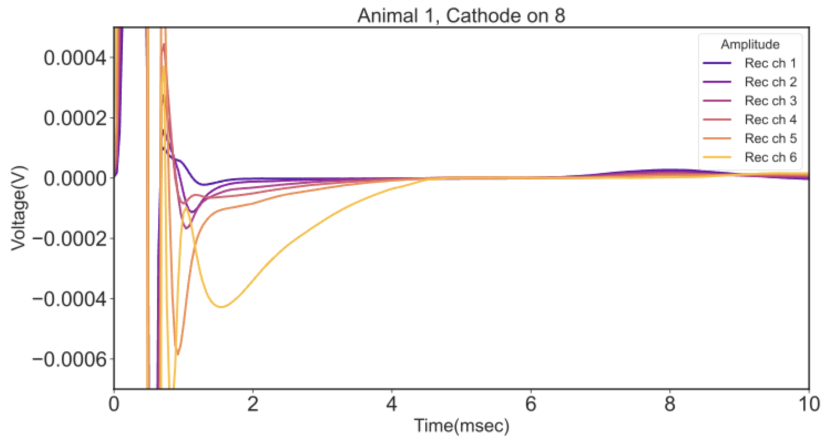


**Animal 1 (cathode on 8):** Vecuronium - Recording Channels 1-6

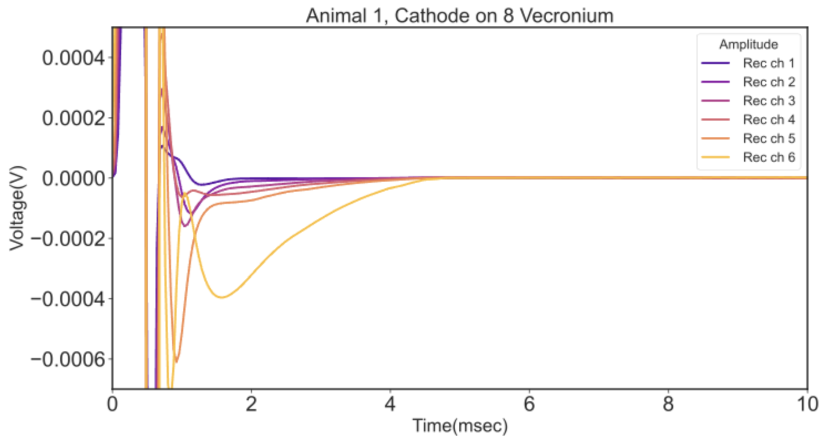


**Animal 1 (cathode on 8):** Baseline - Recording Channels 9-16.

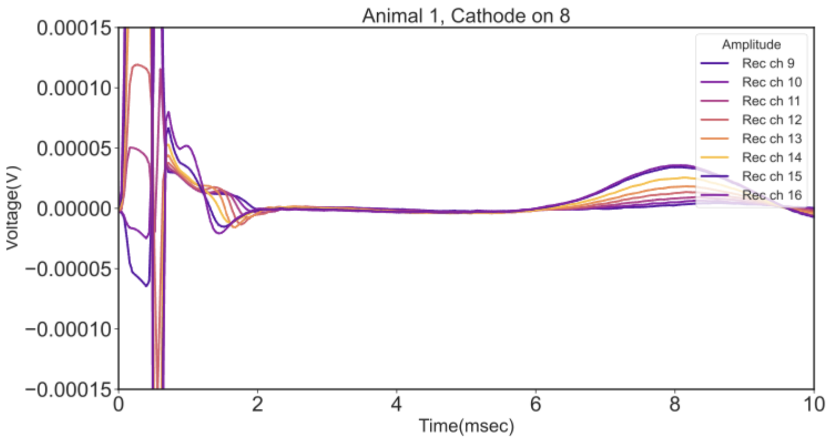


**Animal 1 (cathode on 8):** Vecuronium - Recording Channels 9-16

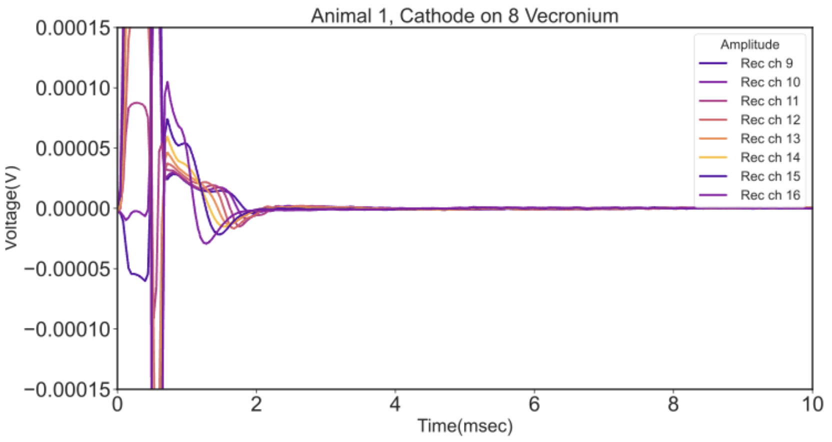

**Animal 2 (cathode on 7):** Baseline - Recording Channels 1-6

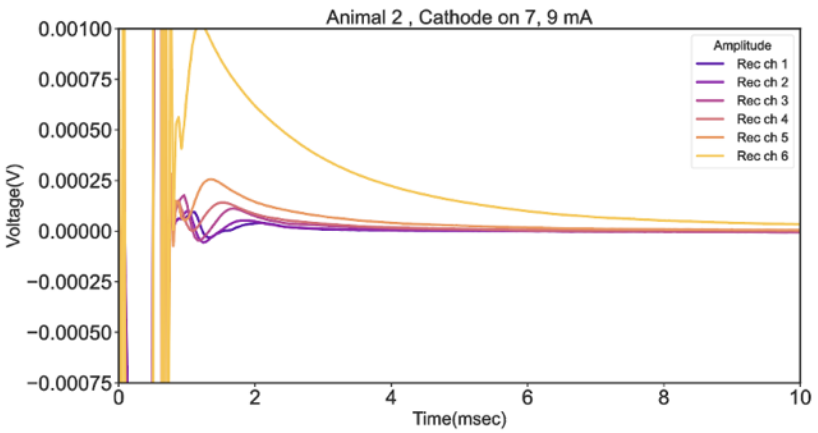


**Animal 2 (cathode on 7):** Vecuronium- Recording Channels 1-6

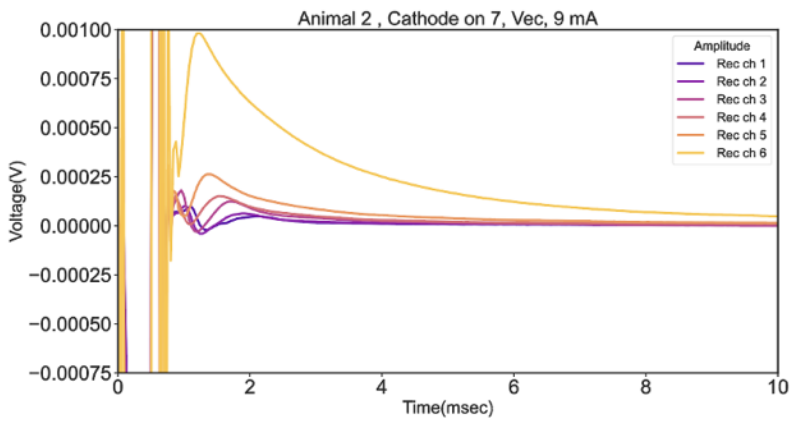


**Animal 2 (cathode on 7):** Baseline - Recording Channels 9-16

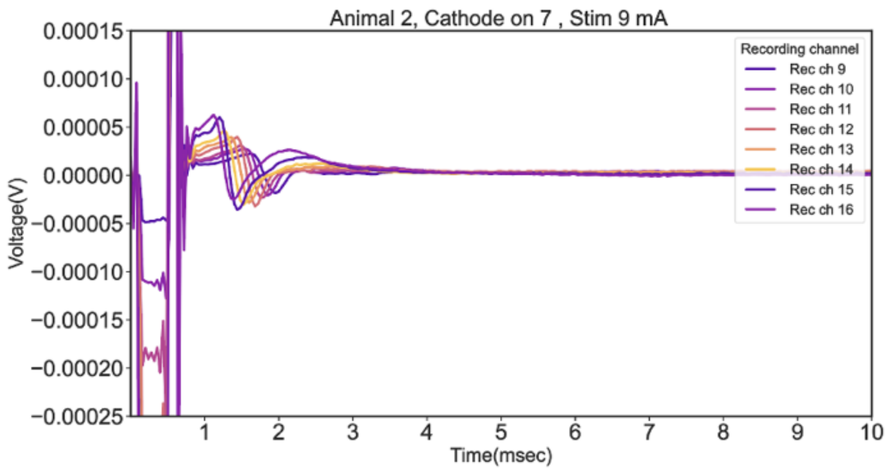


**Animal 2 (cathode on 7):** Vecuronium- Recording Channels 9-16

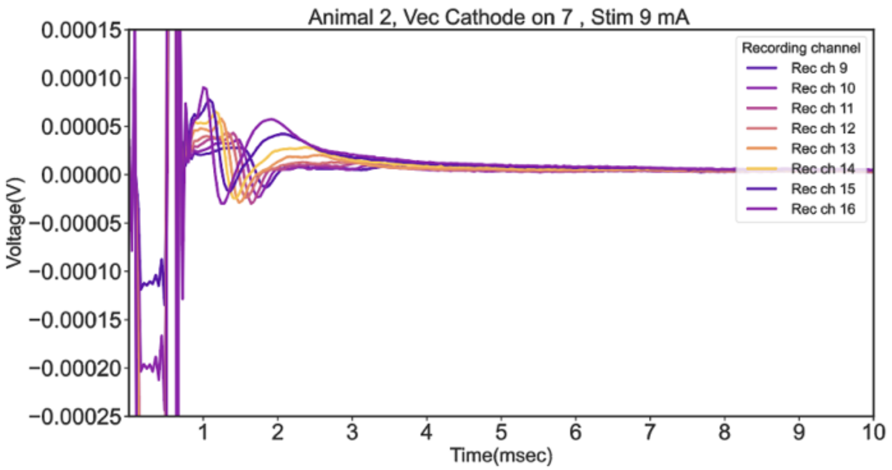

**Animal 2 (cathode on 8):** Baseline - Recording Channels 1-6

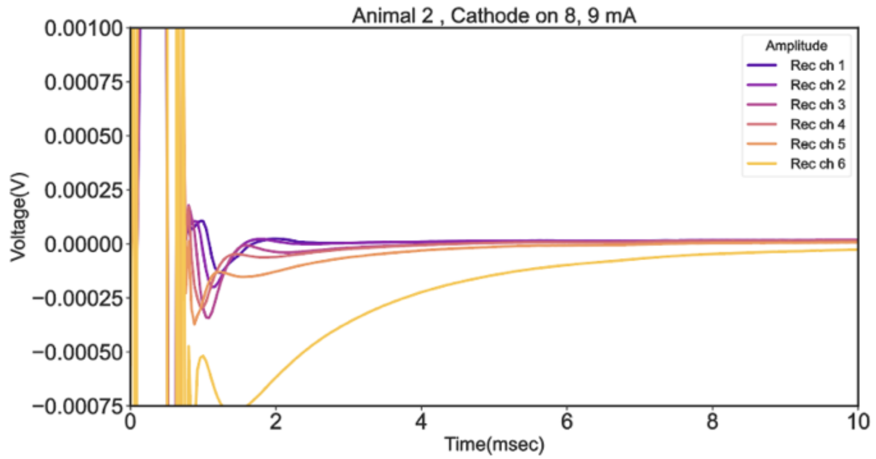


**Animal 2 (cathode on 8):** Vecuronium - Recording Channels 1-6

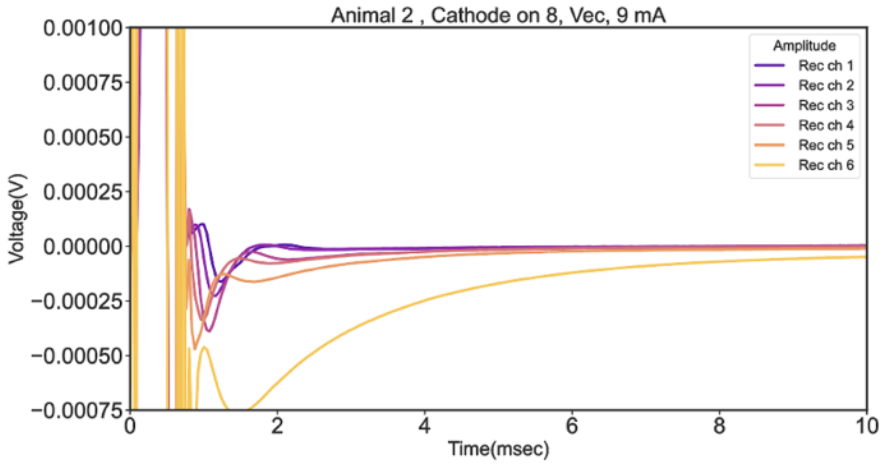


**Animal 2 (cathode on 8):** Baseline - Recording Channels 9-16

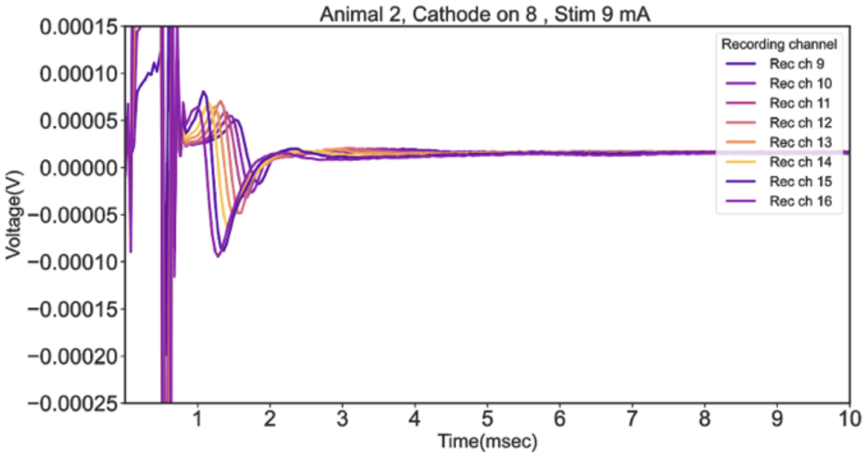


**Animal 2 (cathode on 8):** Vecuronium - Recording Channels 9-16

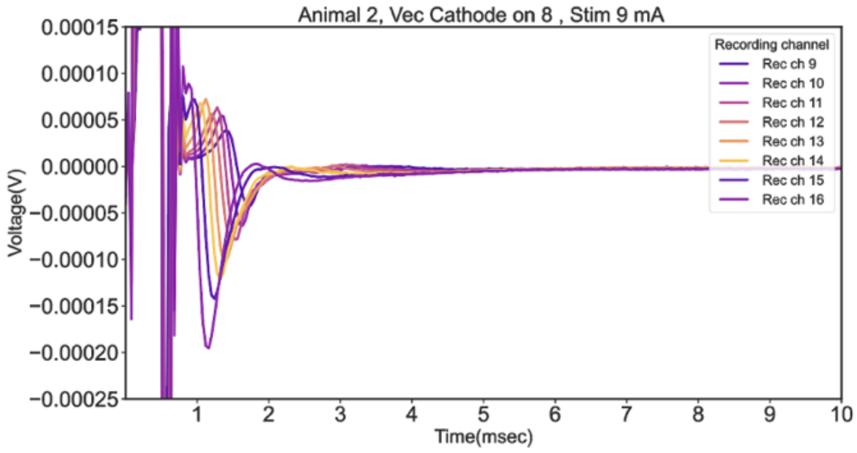

**Animal 4 (cathode on 7):** Baseline - Recording Channels 1-6

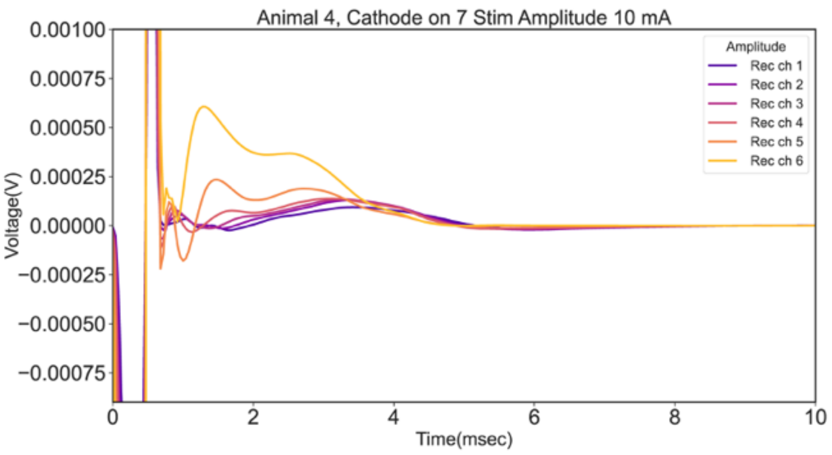


**Animal 4 (cathode on 7):** Vecuronium- Recording Channels 1-6

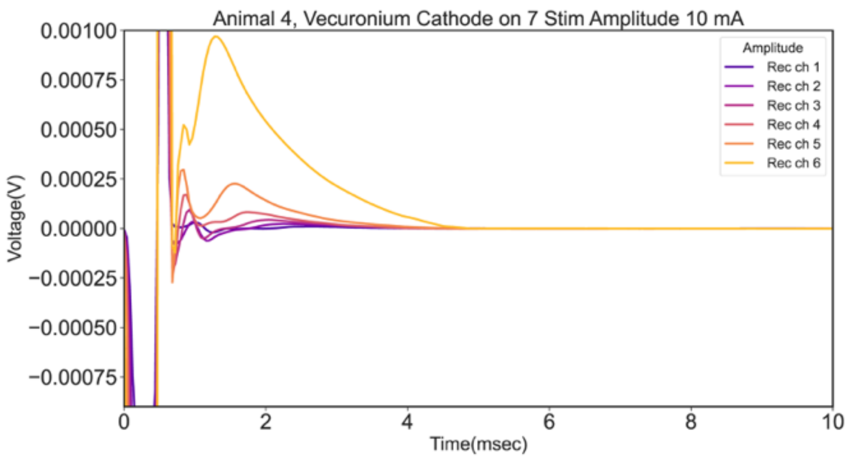


**Animal 4 (cathode on 7):** Baseline - Recording Channels 9-16

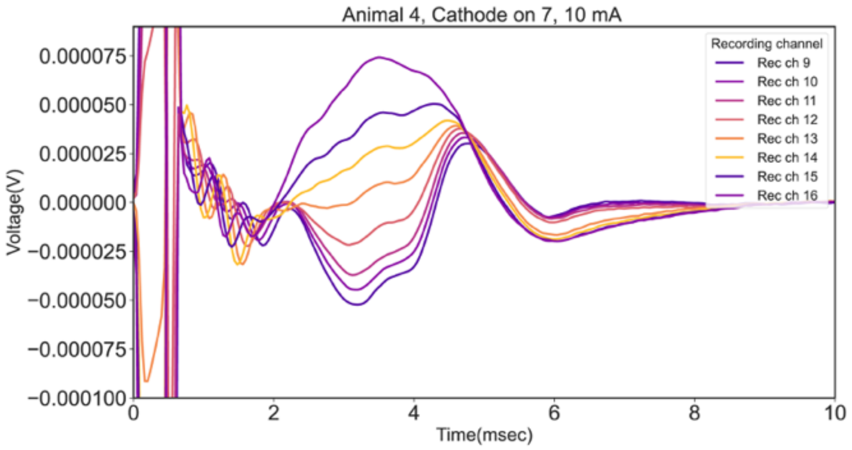


**Animal 4 (cathode on 7):** Vecuronium - Recording Channels 9-16

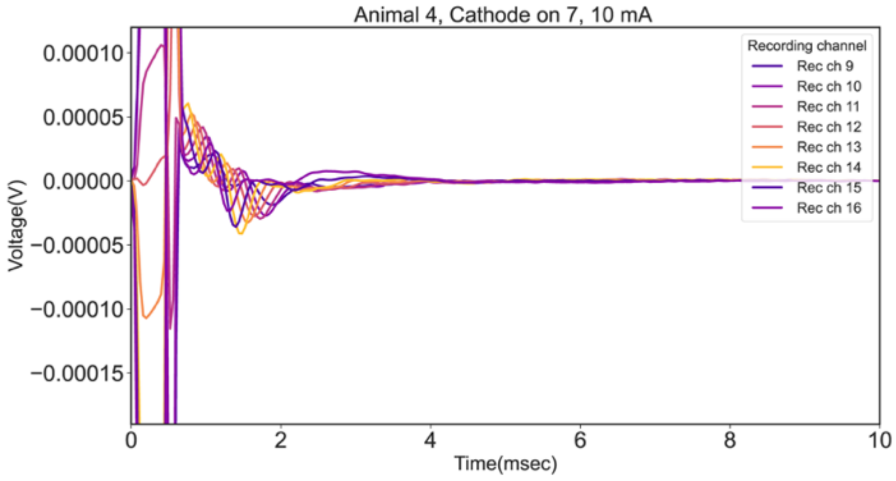

**Animal 4 (cathode on 8):** Baseline - Recording Channels 1-6

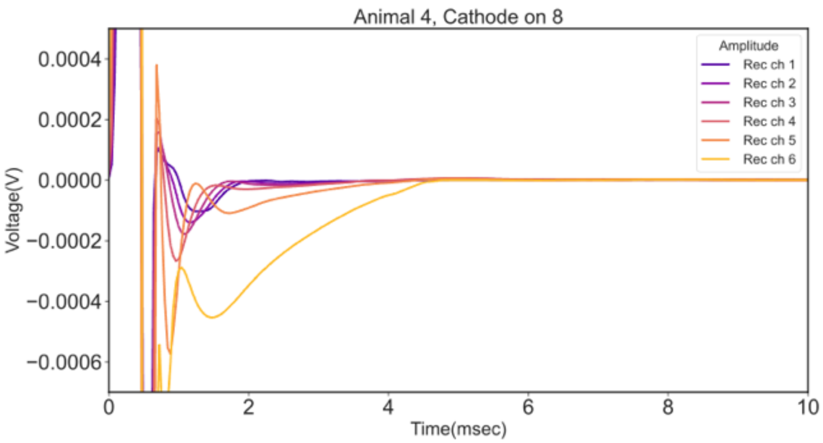


**Animal 4 (cathode on 8):** Vecuronium- Recording Channels 1-6

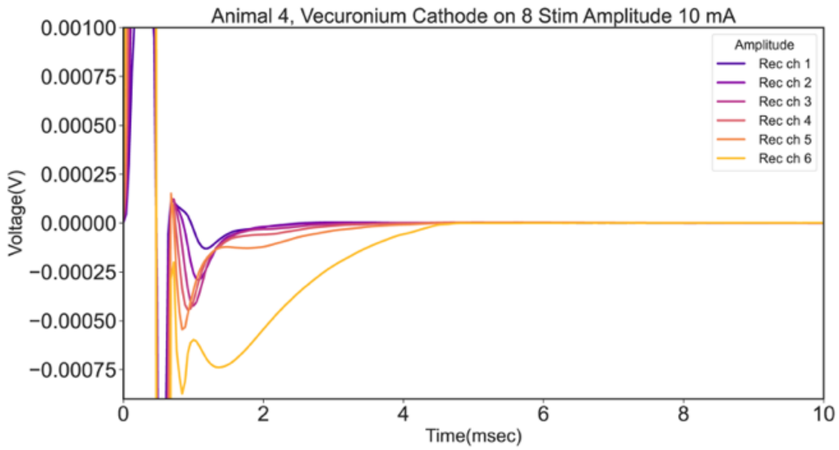


**Animal 4 (cathode on 8):** Baseline - Recording Channels 9-16

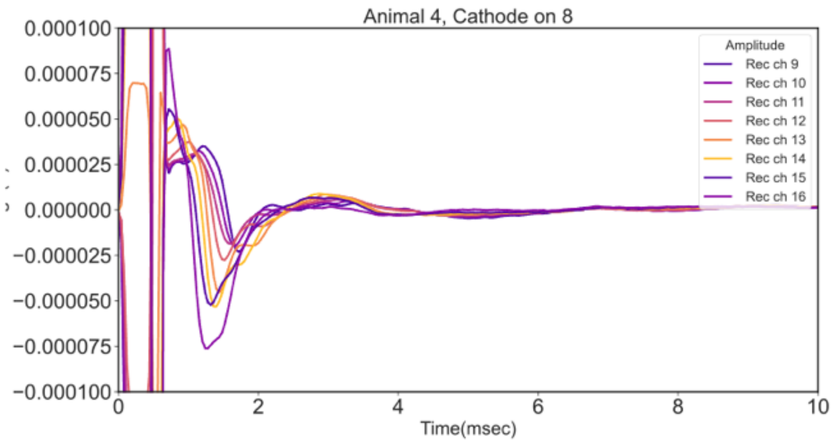


**Animal 4 (cathode on 8):** Vecuronium- Recording Channels 9-16

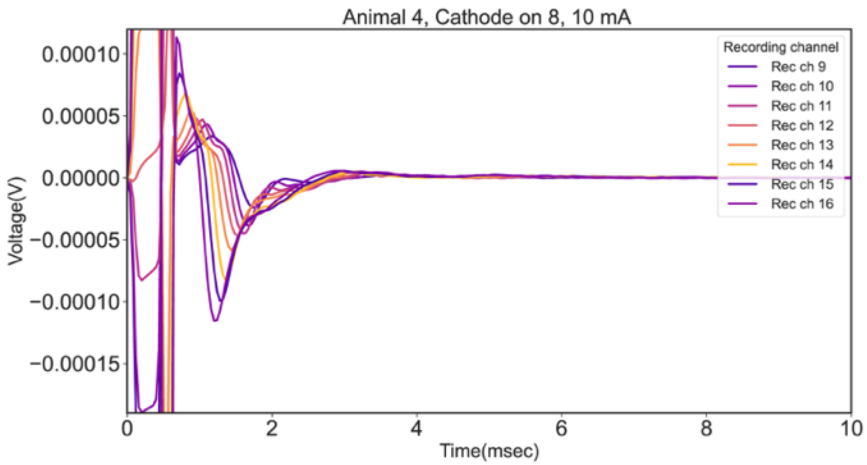

**Animal 5 (cathode at 7):** Baseline - Recording Channels 1-6

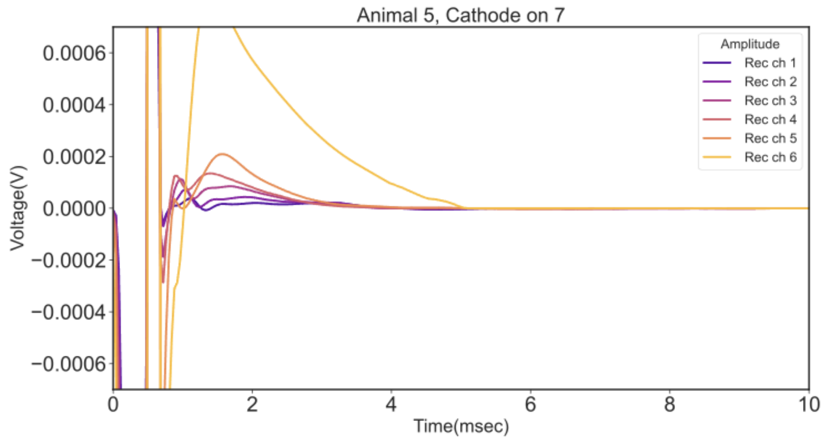


**Animal 5 (cathode at 7):** Vecuronium - Recording Channels 1-6

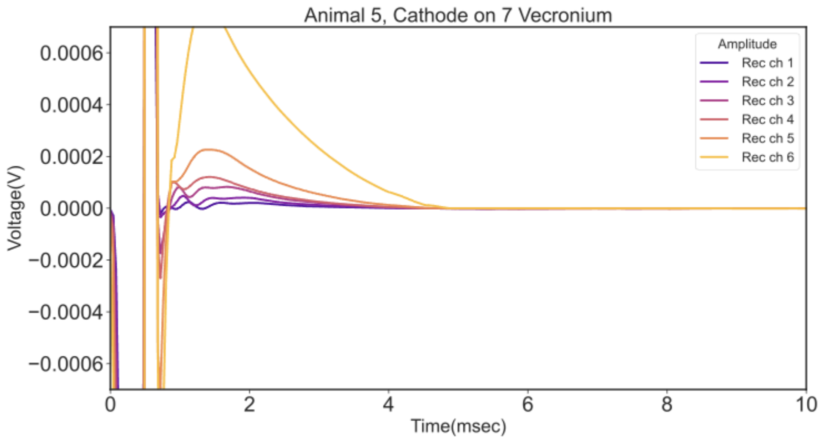


**Animal 5 (cathode at 7):** Baseline - Recording Channels 9-16

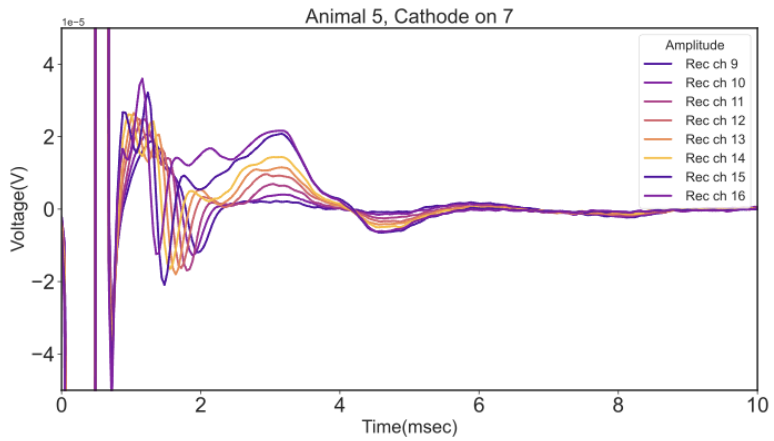


**Animal 5 (cathode at 7):** Vecuronium - Recording Channels 9-16

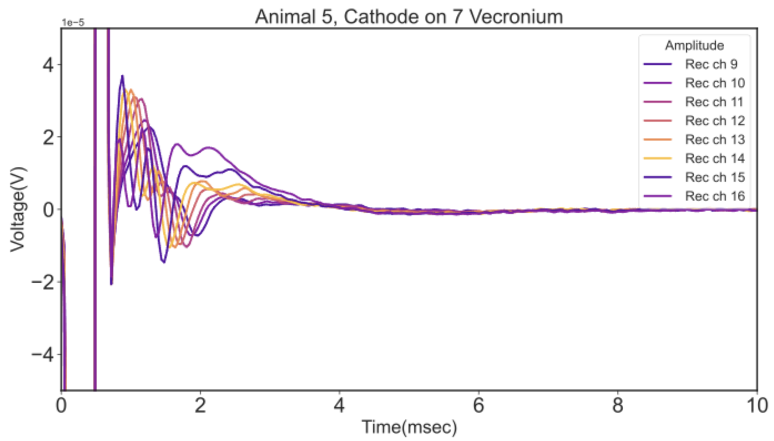

**Animal 5 (cathode at 8):** Baseline - Recording Channels 1-6

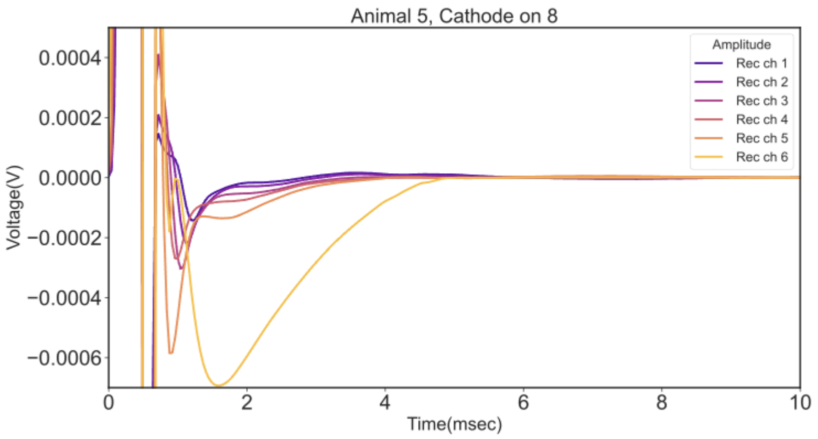


**Animal 5 (cathode at 8):** Vecuronium - Recording Channels 1-6

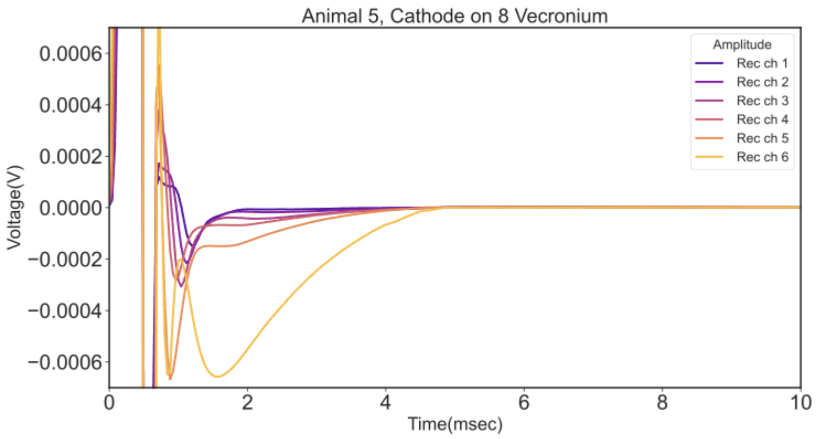


**Animal 5 (cathode at 8):** Baseline - Recording Channels 9-16

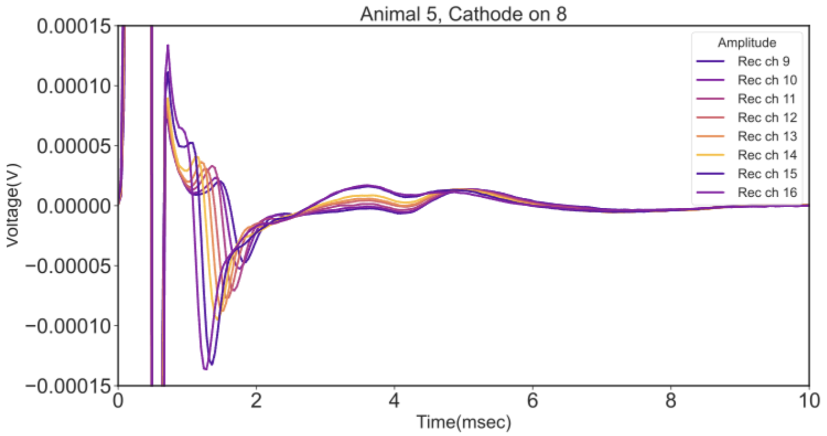


**Animal 5 (cathode at 8):** Vecuronium - Recording Channels 9-16

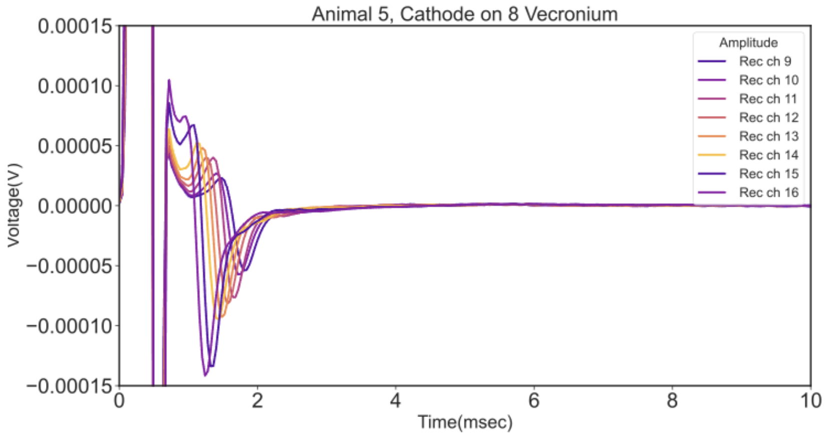

**Animal 6 (cathode at 7):** Baseline - Recording Channels 1-6

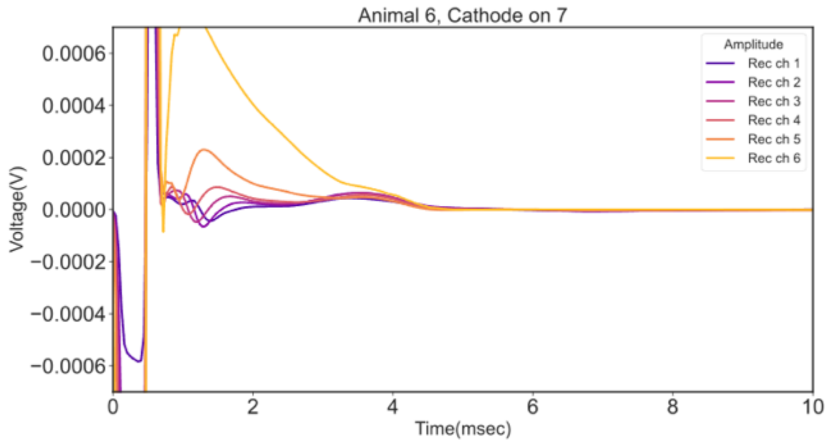


**Animal 6 (cathode at 7):** Vecuronium- Recording Channels 1-6

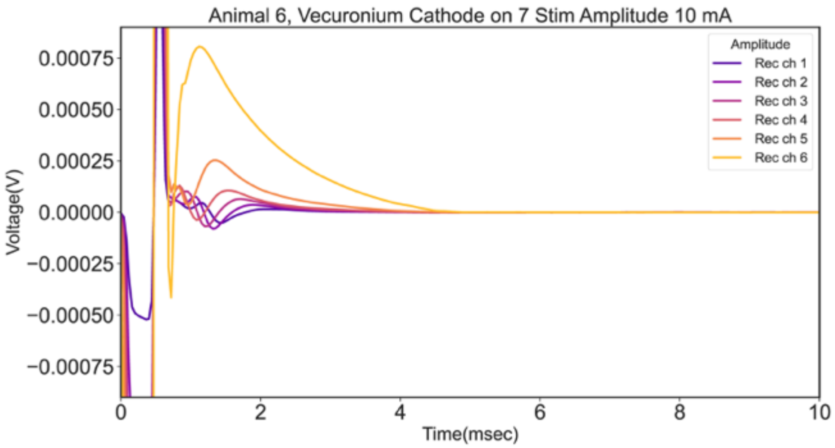


**Animal 6 (cathode at 7):** Baseline - Recording Channels 9-16

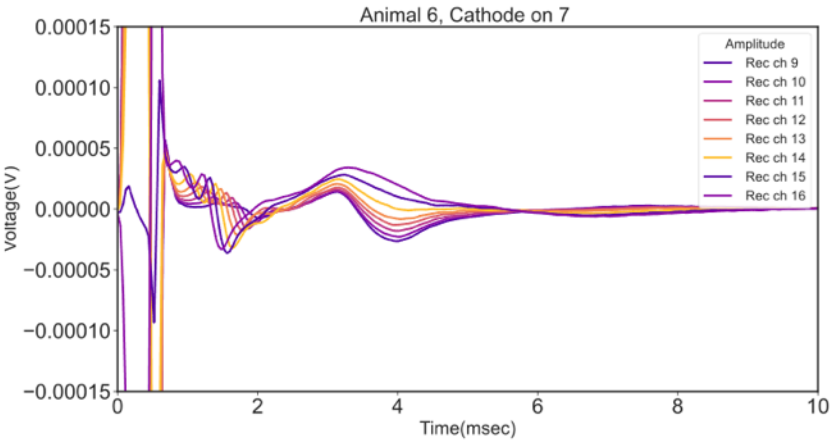


**Animal 6 (cathode at 7):** Vecuronium- Recording Channels 9-16

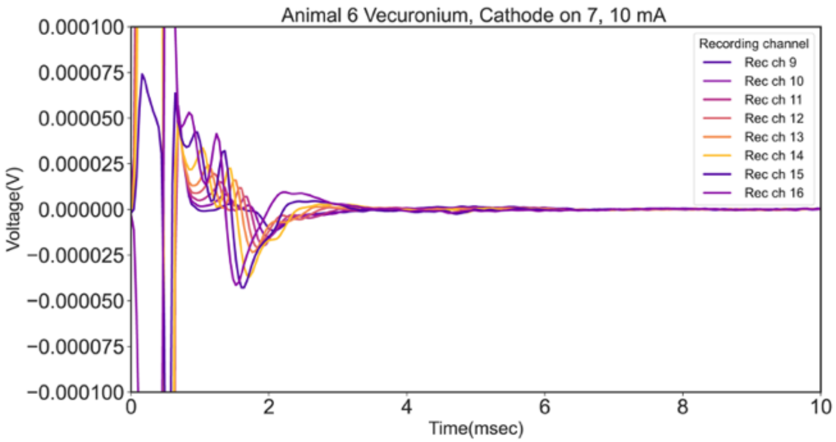

**Animal 6 (cathode at 8):** Baseline - Recording Channels 1-6

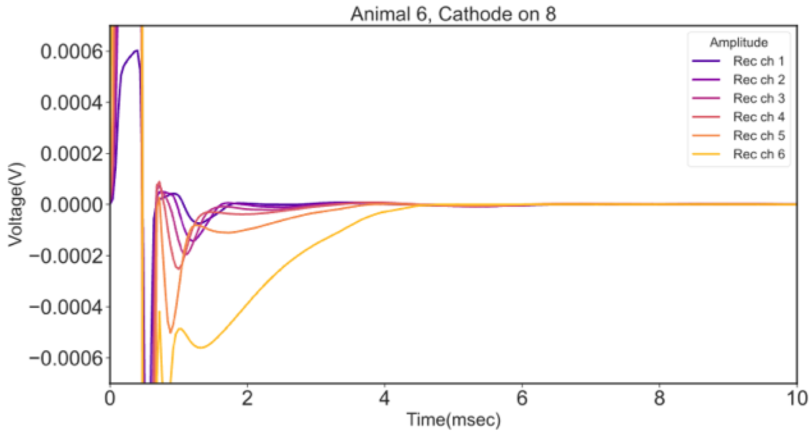


**Animal 6 (cathode at 8):** Vecuronium- Recording Channels 1-6

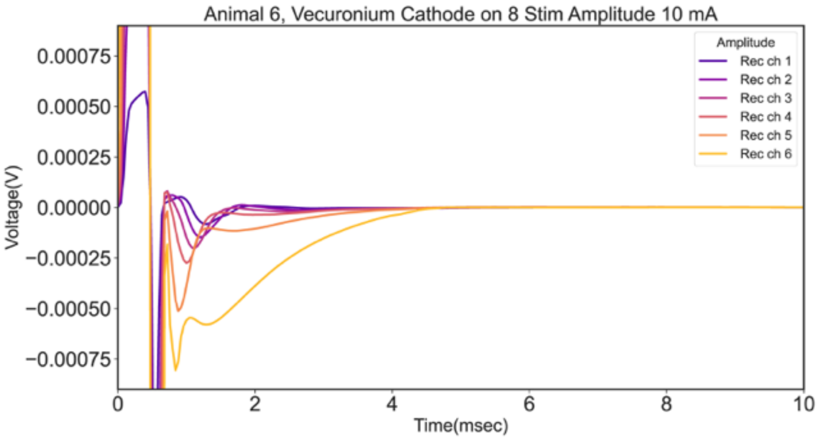


**Animal 6 (cathode at 8):** Baseline - Recording Channels 9-16

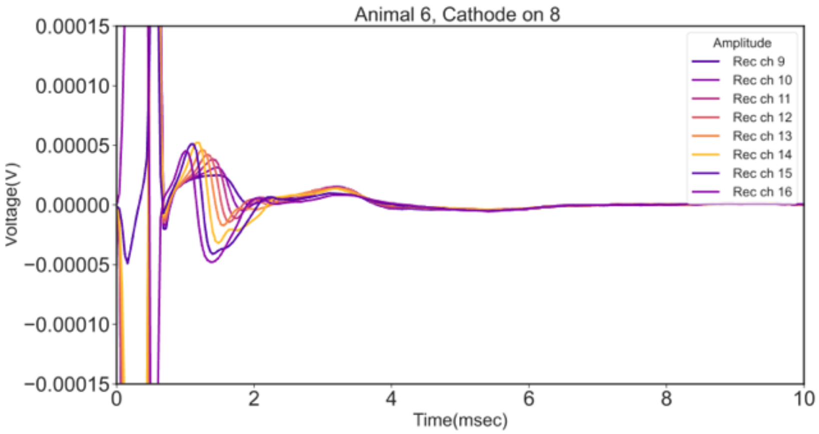


**Animal 6 (cathode at 8):** Vecuronium- Recording Channels 9-16

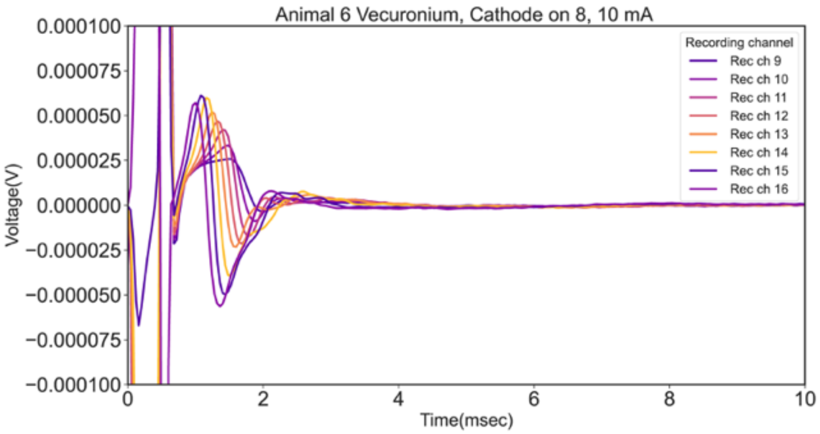

**Animal 7 (cathode at 7):** Baseline - Recording Channels 1-6

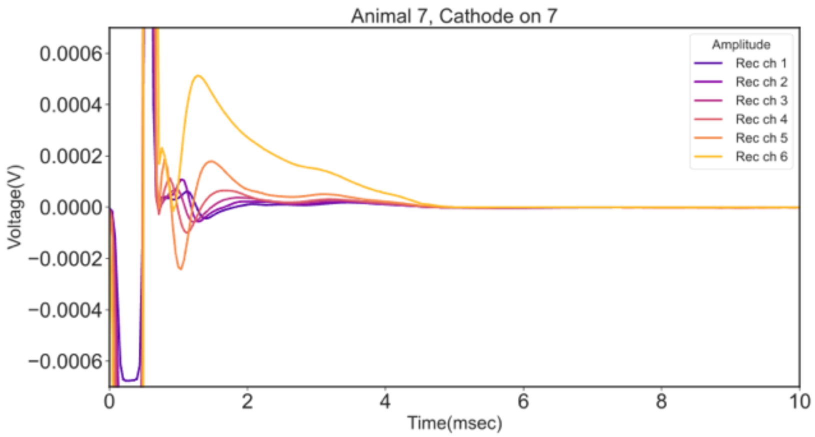


**Animal 7 (cathode at 7):** Vecuronium- Recording Channels 1-6

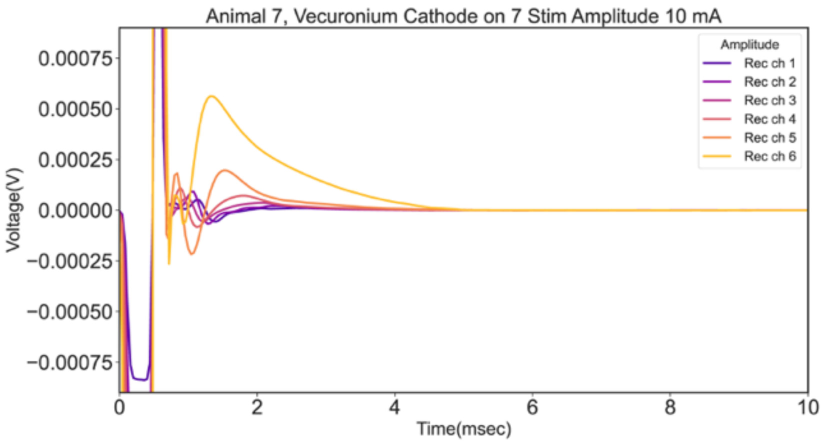


**Animal 7 (cathode at 7):** Baseline - Recording Channels 9-16

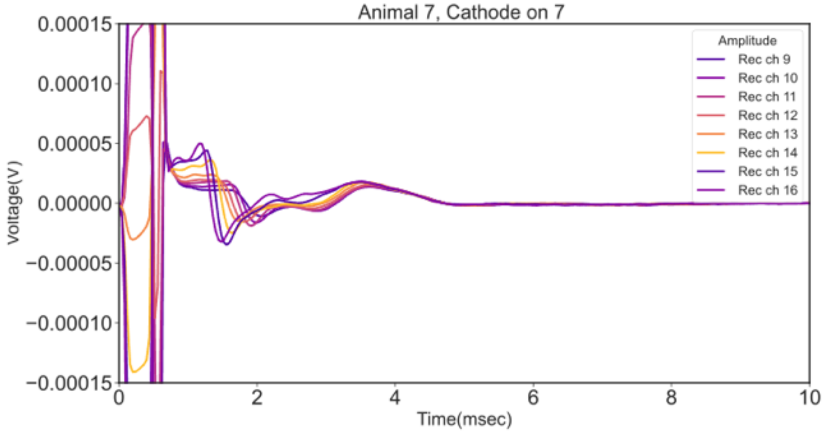


**Animal 7 (cathode at 7):** Vecuronium - Recording Channels 9-16

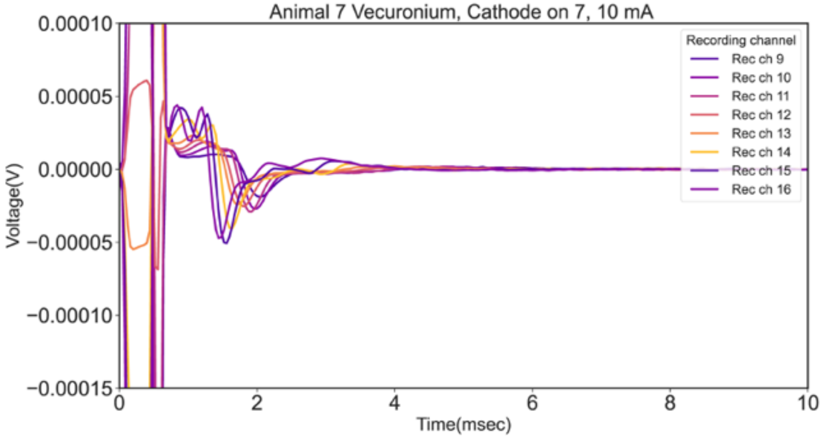

**Animal 7 (cathode at 8):** Baseline - Recording Channels 1-6

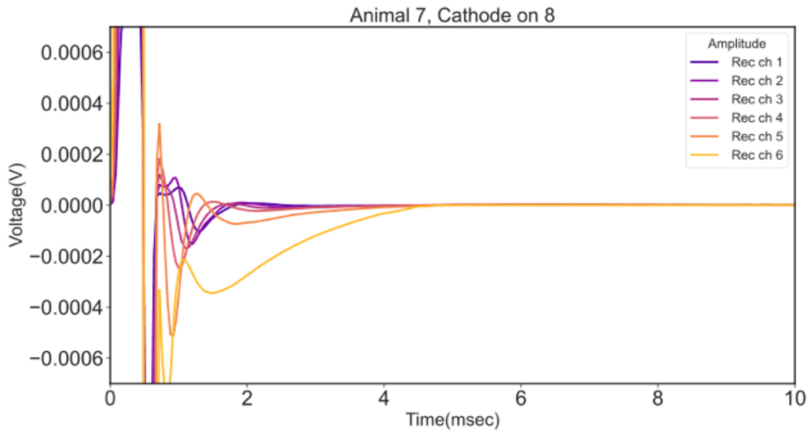


**Animal 7 (cathode at 8):** Vecuronium - Recording Channels 1-6

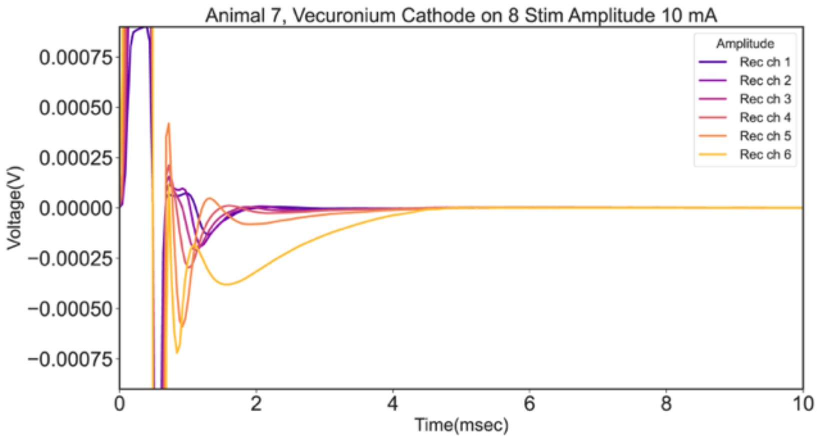


**Animal 7 (cathode at 8):** Baseline - Recording Channels 9-16

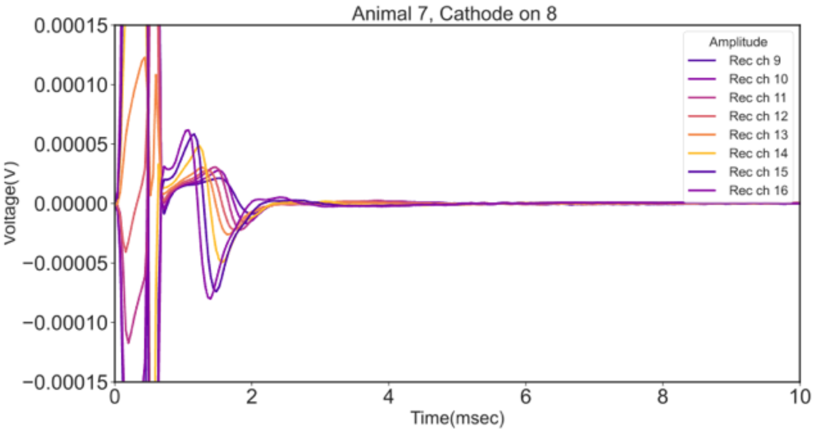


**Animal 7 (cathode at 8):** Vecuronium - Recording Channels 9-16

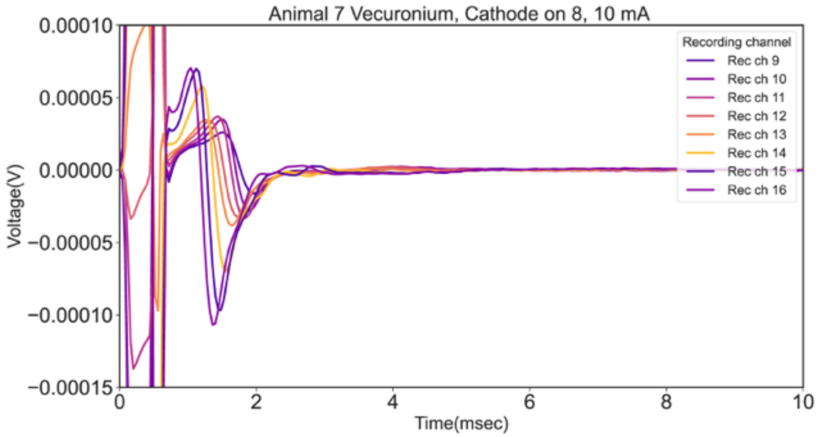

**Supplemental Figure 5:** Post-mortem stimulation-induced capacitive artifact DRC for each subject. Column one is ECAP recordings during stimulation with the cathode on contact 7 (forward polarity), and column two is with stimulation with the cathode on contact 8 (reverse polarity).

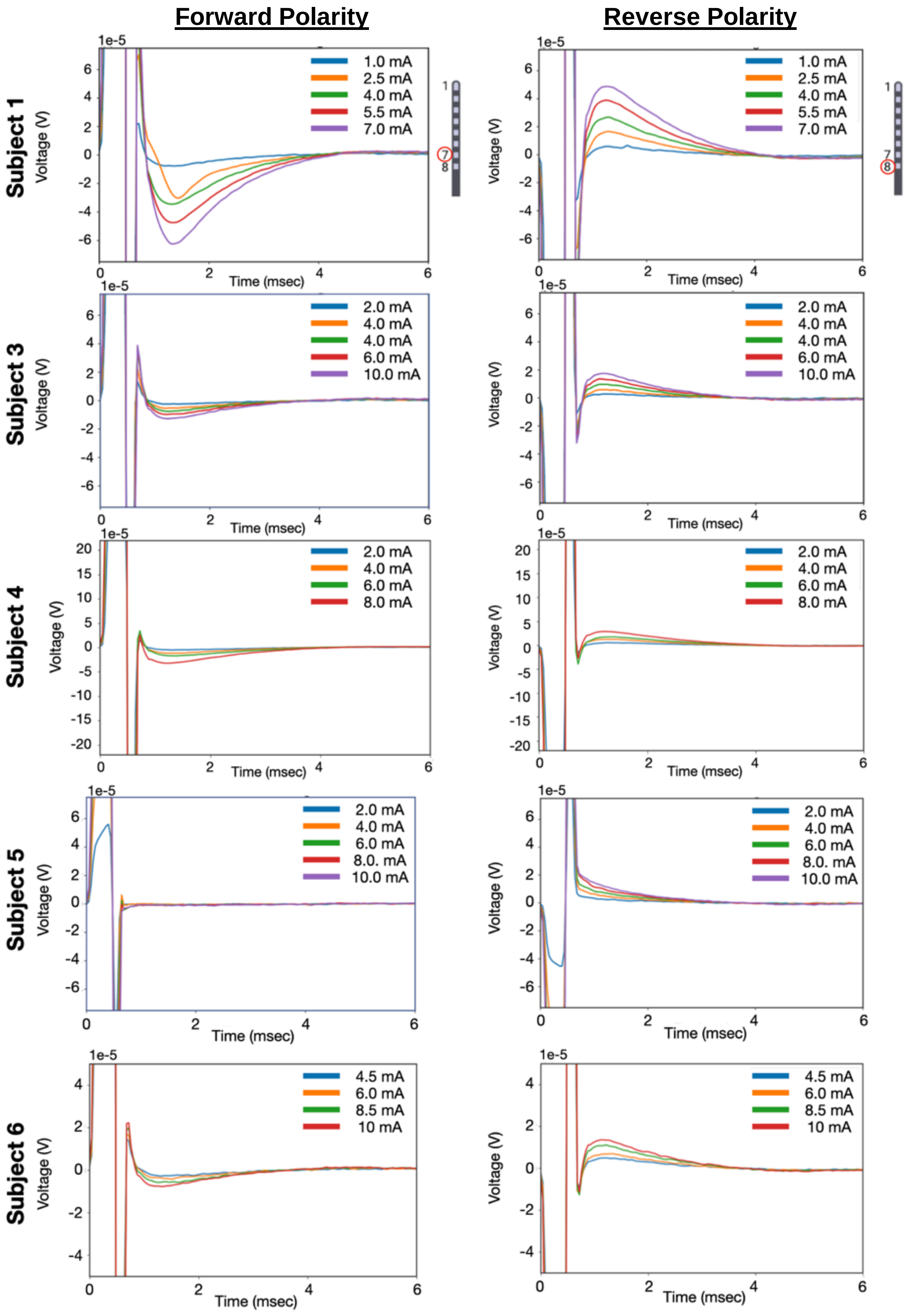

**<u>Supplemental Figure 6:</u>** Representative evolution of double hump with increasing amplitudes across animal 1. The 2nd component was only present in one stimulation configuration with cathode leading asymmetric waveform. The 2nd component was verified to be neural by carrying out the analysis under Vecuronium. The green dotted lines show the peaks marked to calculate the latency for each animal (Latency Bar Graph, below). The conduction velocity calculation is also shown for representative animal 1 (Conduction Velocity Bar Graph, below).

**Animal 1:**

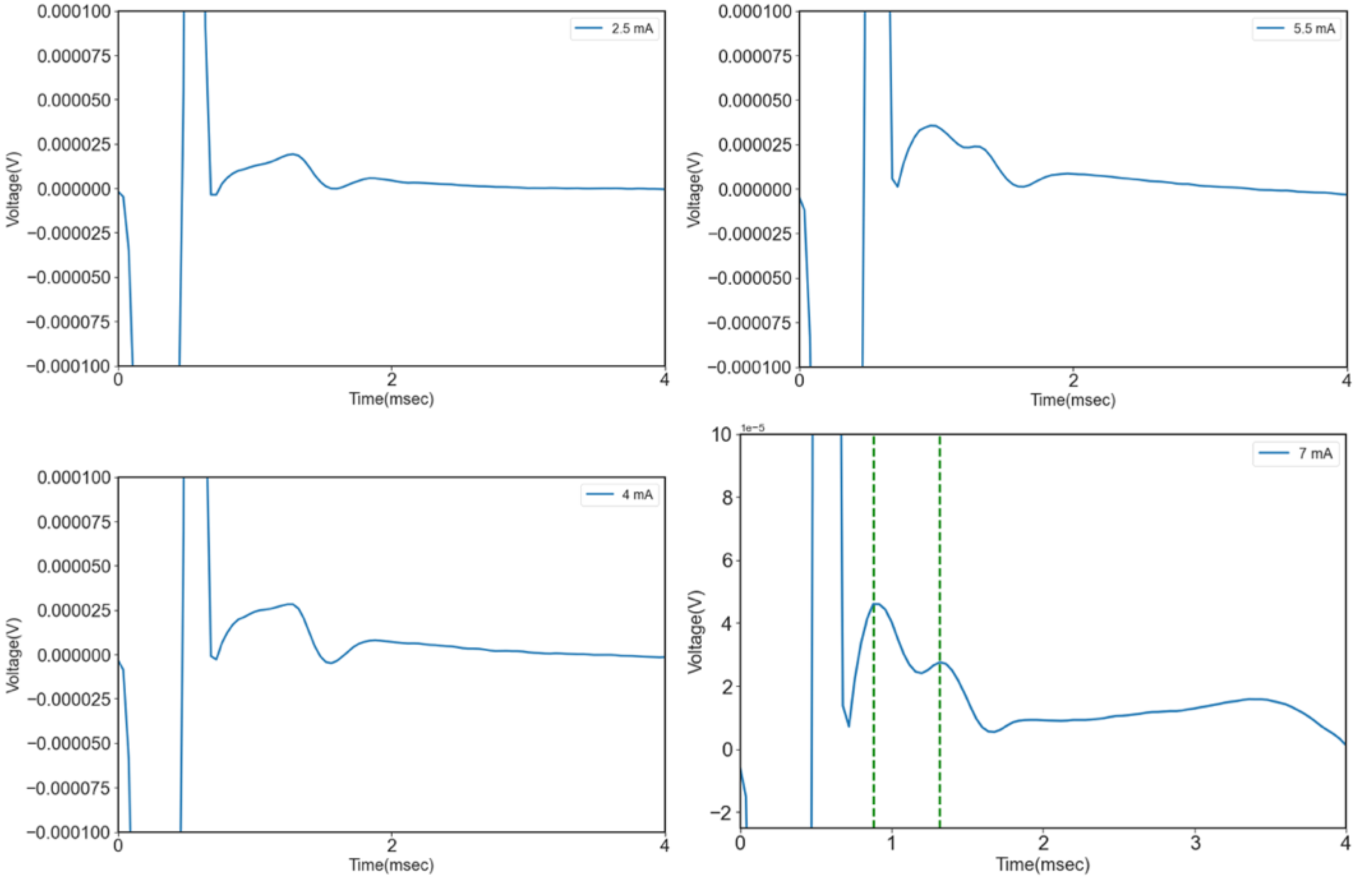


- Approximate distance between direct dorsal column and indirect activation pathway of roots: 2 cm or 20mm
- Time difference between two peaks = 0.44msec
- Conduction velocity for peak : 20/0.44 = 45 m/sec

**Supplementary Figure 6:**

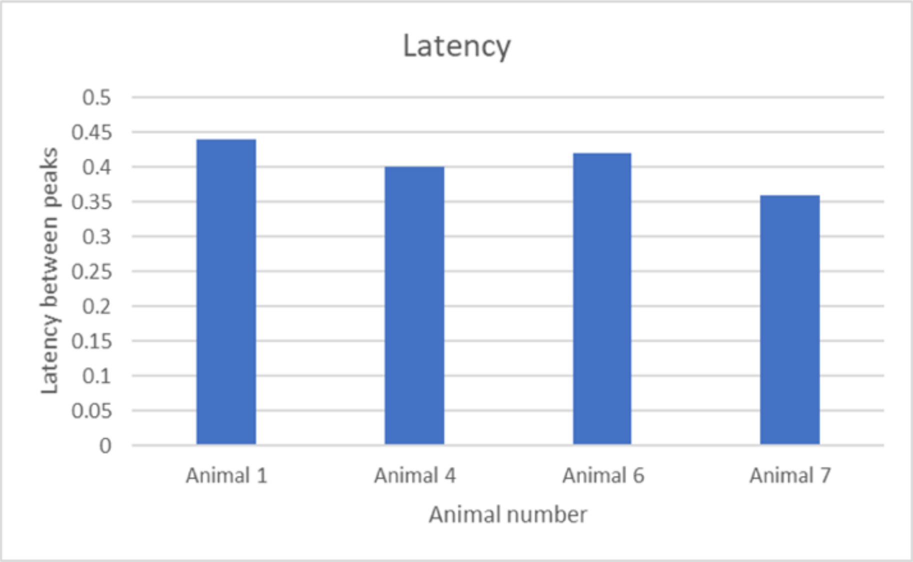


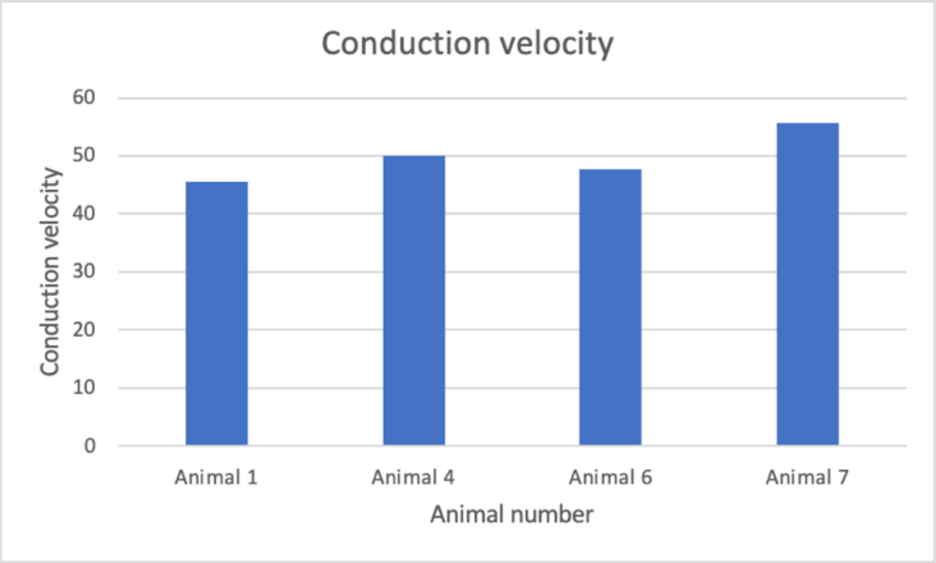


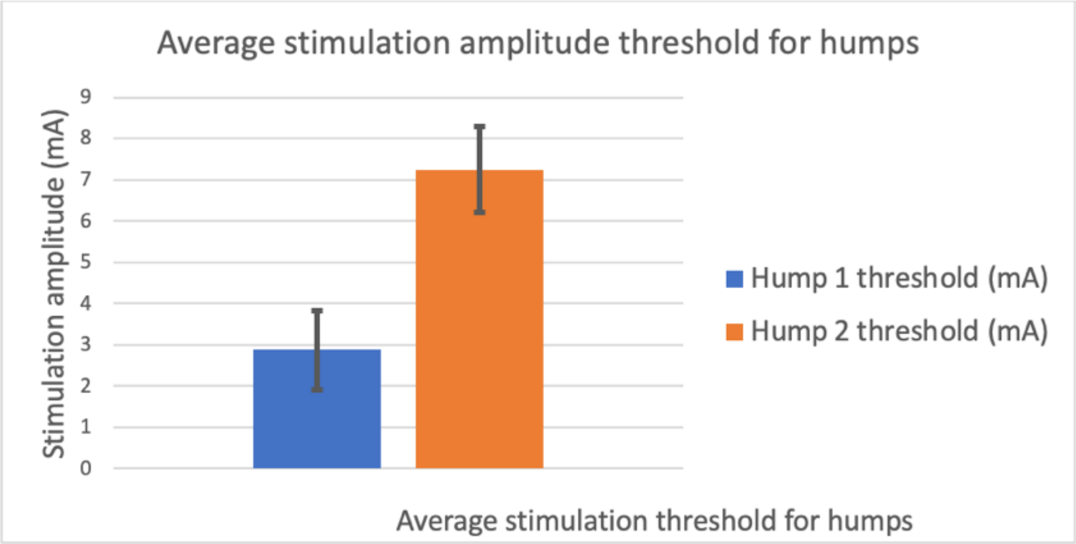

**Supplemental Figure 7:** Double hump data for all animals under Vecuronium. Comparing cathode at Channel (ch) 7 vs ch 8.

**Animal 1:**

Anode ch 8, cathode ch 7

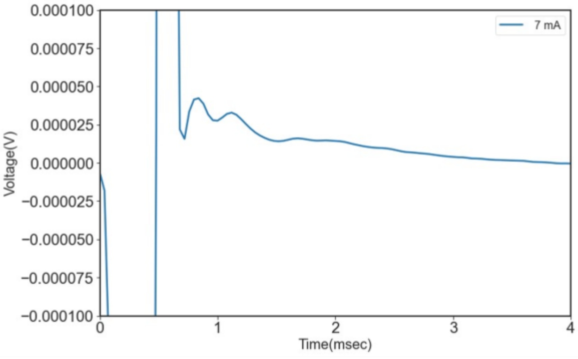


Anode ch 7, cathode ch 8

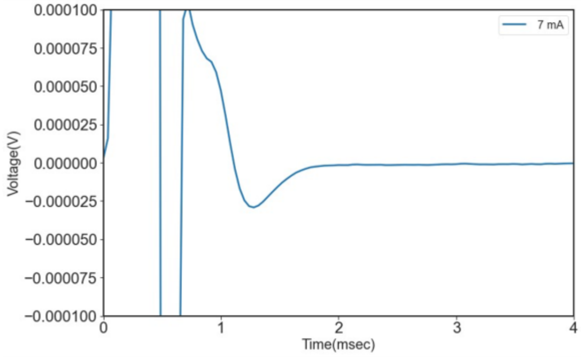


**Animal 2:**

Anode ch 8, cathode ch 7

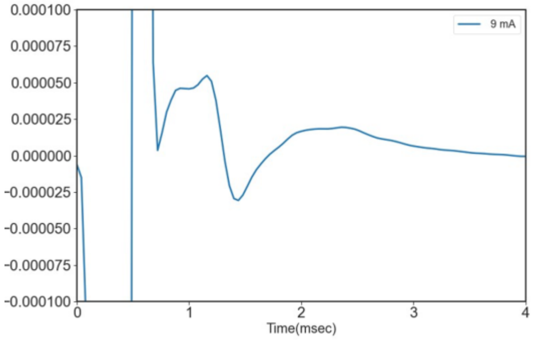


Anode ch 7, cathode ch 8

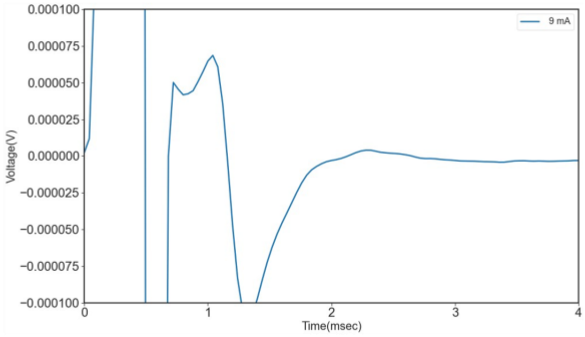


**Animal 3**:

Anode ch 8, cathode ch 7

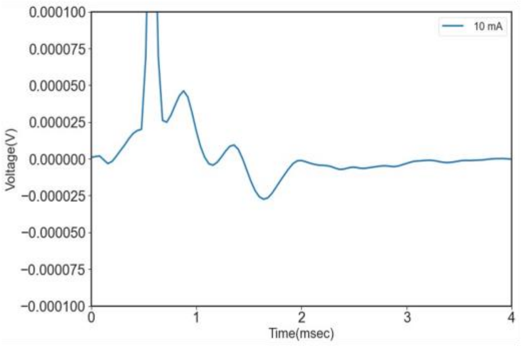


Anode ch 7, cathode ch 8

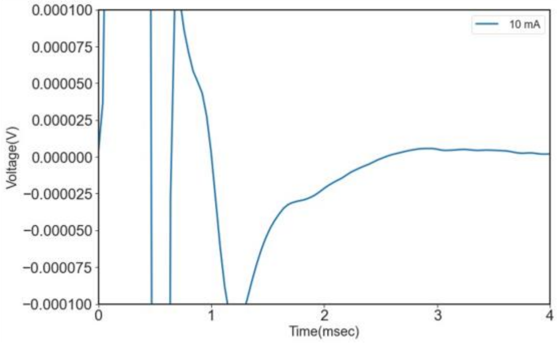

**Animal 4:**

Anode ch 8, cathode ch 7

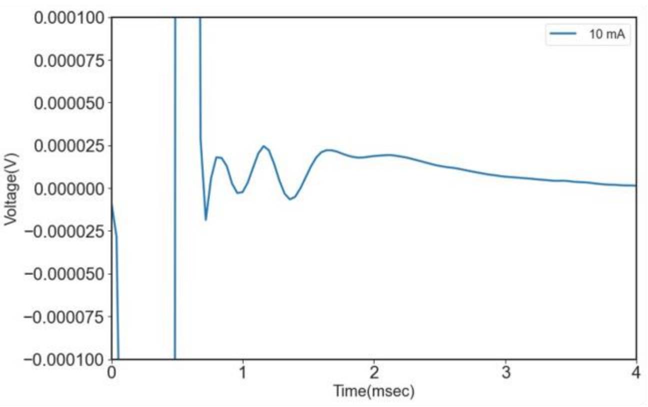


Anode ch 7, cathode ch 8

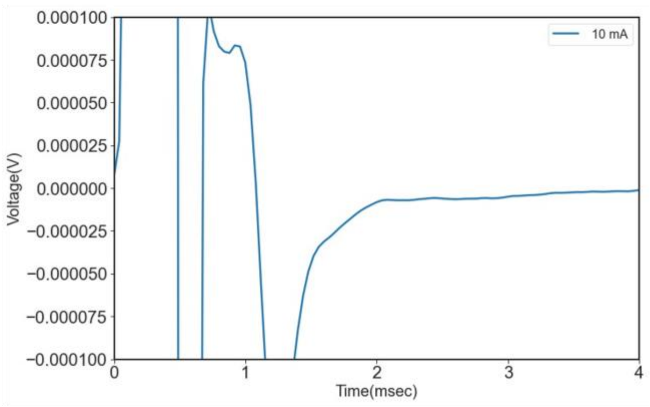


**Animal 5:**

Anode ch 8, cathode ch 7

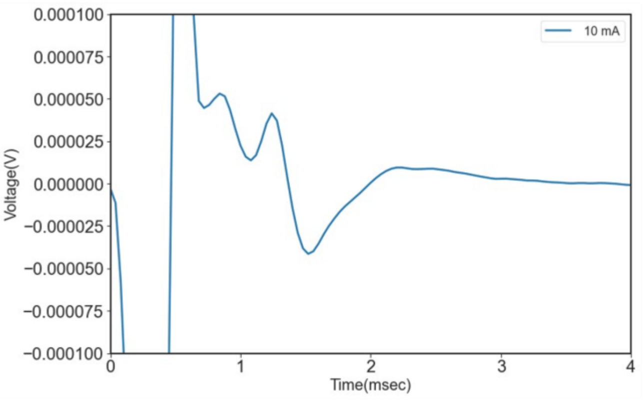


Anode ch 7, cathode ch 8

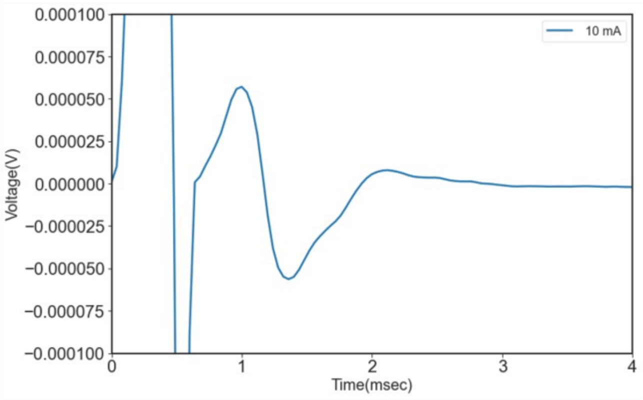


**Animal 6:**

Anode ch 8, cathode ch 7

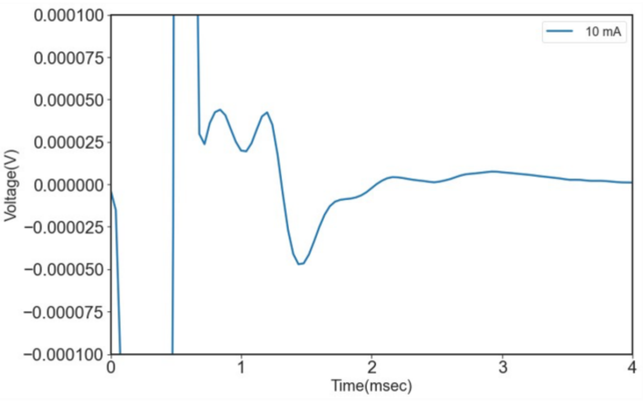


Anode ch 7, cathode ch 8

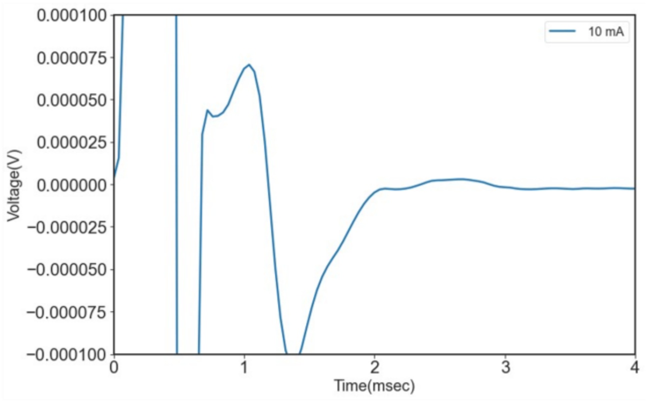

**<u>Supplemental Figure 8:</u>** Two representative examples of transient artifacts during spinal cord ECAP recording (black arrows) in subject 2 (left) and subject 4 (right).

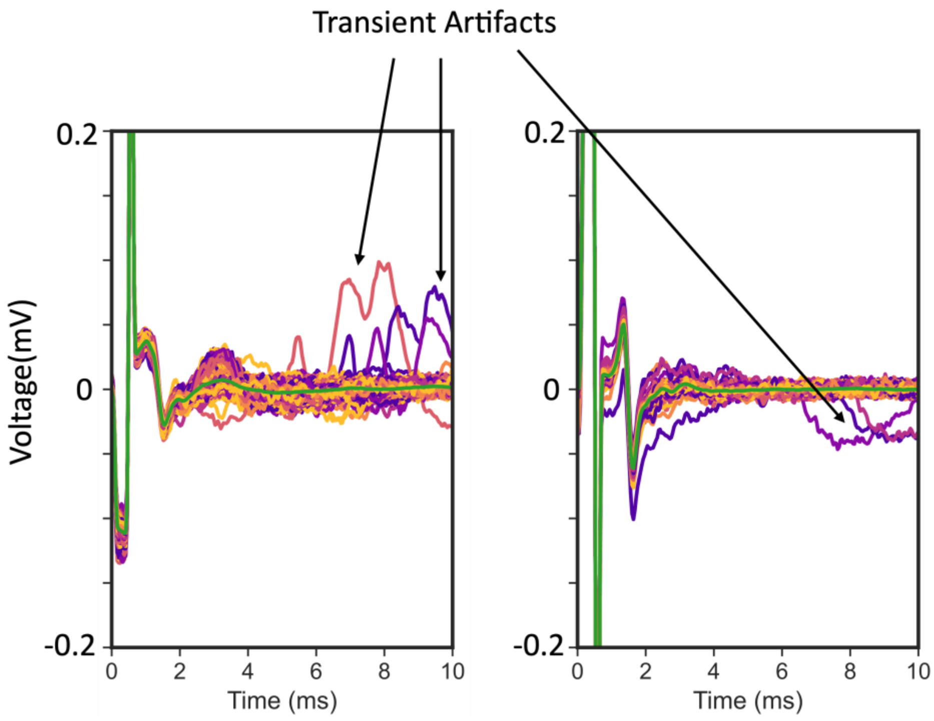
